# High level and molecular nature of transcriptional noise in yeast cells

**DOI:** 10.1101/2022.10.26.513925

**Authors:** Zlata Gvozdenov, Zeno Barcutean, Kevin Struhl

## Abstract

“Biological noise” is defined as functionally insignificant events that occur in living cells due to imperfect fidelity of biological processes. Distinguishing between biological function and biological noise is often difficult, and experiments to measure biological noise have not been performed. Here, we measure biological noise in yeast cells by analyzing chromatin structure and transcription of an 18 kb region of DNA whose sequence was randomly generated and hence lacks biological function. Nucleosome occupancy on random-sequence DNA is comparable to that on yeast genomic DNA. However, nucleosome-depleted regions are much less frequent, and there are fewer well-positioned nucleosomes and shorter nucleosome arrays. Steady-state levels of RNAs expressed from random-sequence DNA are comparable to those of typical yeast mRNAs, although transcription and mRNA decay rates are higher. Transcriptional initiation (5’ ends) from random-sequence DNA occurs at numerous sites at low levels, indicating very low intrinsic specificity of the Pol II machinery. In contrast, poly(A) profiles (relative levels and clustering of 3’ isoforms) of random-sequence RNAs are roughly comparable to those within 3’ untranslated regions of yeast mRNAs, suggesting limited evolutionary constraints on poly(A) site choice. RNAs expressed from random-sequence DNA show higher cell-to-cell variability than RNAs expressed from yeast genomic DNA, suggesting that functional elements limit the variability among individual cells within a population. These observations indicate that transcriptional noise occurs at high levels in yeast, and they provide insight into how chromatin and transcription patterns arise from the evolved yeast genome.

## INTRODUCTION

Biological function, while often obvious, is an elusive concept. Are all events that occur in living cells biologically meaningful? The flip side of biological function is a concept termed “biological noise”, which is defined by reproducible events (not experimental error) that occur in living cells due to imperfection (i.e., lack of fidelity) of a biological process (Struhl 2007). Biological noise is distinct from stochastic variations that occur in individual cells.

The classic view of transcription by RNA polymerase II (Pol II) is that initiation from the promoter and cleavage/polyadenylation in the 3’ untranslated region (3’ UTR) leads to discrete and functional RNA species, most of which are mRNAs. There is considerable heterogeneity at 5’ and 3’ ends of these mRNAs, leading to hundreds of different isoforms (Pelechano et al. 2013). However, there are numerous Pol II-generated transcripts that are not initiated from canonical promoters, do not terminate at the termination regions, and can be transcribed in the opposite direction of classic mRNAs (David et al. 2006; Neil et al. 2009; Xu et al. 2009). Many “cryptic” RNAs are unstable in wild-type cells, and they are only observed in strains defective for genes involved in RNA degradation (Wyers et al. 2005; Arigo et al. 2006; Thiebaut et al. 2006) or in maintaining repressive chromatin structure (Kaplan et al. 2003; Mason and Struhl 2003; Carrozza et al. 2005; Joshi and Struhl 2005; Keogh et al. 2005). Are these non-classical RNAs functional or do they represent biological noise?

Based on the genome-wide distribution of Pol II and quantitative information in yeast cells, it was estimated that only ∼10% of the ∼12,000 elongating molecules at any given time are engaged in the generation of conventional mRNAs (Struhl 2007). Calculations of Pol II specificity suggest a 10^4^-fold difference between an optimal promoter vs. a random genomic site. Interestingly, this level of specificity is comparable to that of a typical sequence-specific DNA-binding protein and to the frequency at which incorrect nucleotides or amino acids are incorporated into RNA and protein polymers. From these observations, it was proposed that ∼90% of elongating Pol II molecules represent transcriptional noise, producing transcription with no biological role (Struhl 2007). However, transcriptional noise has never been experimentally defined or measured.

Pol II transcription is inextricably intertwined with chromatin structure. As Pol II traverses the gene, there is a rapid equilibrium between histone eviction to facilitate Pol II elongation and histone deposition to restore normal chromatin structure (Kristjuhan and Svejstrup 2004; Schwabish and Struhl 2004; Lee et al. 2007). This dynamic process is mediated by histone chaperones and chromatin-modifying proteins such as FACT, Spt6, Asf1, the Rpd3-S histone deacetylase complex, Set2 histone methylase, and the Swi/Snf nucleosome remodeling complex (Kaplan et al. 2003; Mason and Struhl 2003; Carrozza et al. 2005; Joshi and Struhl 2005; Keogh et al. 2005; Schwabish and Struhl 2006; Schwabish and Struhl 2007). Many of these proteins travel with Pol II, either via direct or indirect interactions, or by recognizing the altered structure generated by Pol II transcription. Mutations that inhibit histone deposition result in aberrant open chromatin regions that generate “cryptic” transcripts that are observed at much lower levels in wild-type strains. Genes that are very highly transcribed have reduced levels of nucleosome occupancy in the coding region (Iyer and Struhl 1995b; Kristjuhan and Svejstrup 2004; Schwabish and Struhl 2004; Sekinger et al. 2005; Lee et al. 2007).

Eukaryotic promoters that drive transcription of conventional genes are typically depleted of nucleosomes, thereby permitting increased access to transcription factors (Iyer and Struhl 1995b; Sekinger et al. 2005; Yuan et al. 2005; Mavrich et al. 2008; Struhl and Segal 2013). These nucleosome-depleted regions are flanked by highly positioned +1 (downstream) and −1 (upstream) nucleosomes. Nucleosomes in the mRNA coding region are statistically positioned from the +1 nucleosome, with the degree of positioning decreasing progressively at more downstream locations. The position of the +1 nucleosome and the statistically positioned nucleosomes downstream are, respectively, linked to transcriptional initiation and elongation (Struhl and Segal 2013).

The characteristic chromatin pattern is also observed when nucleosome-depleted regions are fortuitously generated in coding regions of foreign yeast DNA sequences present in *S. cerevisiae* (Hughes et al. 2012). Moreover, these fortuitous nucleosome-depleted regions act as bidirectional promoters that give rise to equal levels of transcription in both directions (Jin et al. 2017). Similarly, enhancers located far from mRNA coding regions are also associated with nucleosome-depleted regions and equal levels of bidirectional transcription (Kim et al. 2010; Core et al. 2014; Jin et al. 2017). Taken together, these considerations indicate that chromatin structure will influence both the amount and location of transcriptional noise.

Biological function has mechanistic and evolutionary components, and these are difficult to disentangle by analysis of wild-type cells. In previous work, we developed a functional evolutionary approach to address mechanistic questions *in vivo* independent of evolutionary history (Hughes et al. 2012). This approach involves the use of standard functional assays to measure molecular events on evolutionarily irrelevant DNA. We analyzed chromatin, transcription, and polyadenylation in *S. cerevisiae* cells containing large (200 kb) segments of DNA from other yeast species of (*K. lactis* and the more evolutionary distant *D. hansenii*)(Hughes et al. 2012; Moqtaderi et al. 2013; Jin et al. 2017). However, foreign yeast DNA, although evolutionarily irrelevant in *S. cerevisiae*, still arose through evolutionary selection.

The best experimental definition of biological noise involves molecular events that occur on random-sequence DNA. Here, we address transcriptional noise by extending the functional evolutionary approach to analyze an 18 kb segment of DNA whose sequence was randomly generated. We analyze chromatin structure (nucleosome positioning and occupancy by MNase-seq), transcription (RNA-seq), 3’-end formation by cleavage/polyadenylation (3’READS), and 5’end formation (5’ ends). Our results indicate that 1) chromatin forms efficiently on random-sequence DNA but with fewer nucleosome-depleted regions and with a lower degree of nucleosome positioning, 2) transcriptional noise occurs at levels comparable to that of typical mRNAs, 3) 5’ ends occur with very little specificity, and 4) poly(A) sites of random-sequence RNAs have features in common with natural mRNAs, although they are somewhat more heterogeneous.

## RESULTS

### Generating a yeast strain with a new chromosome containing random-sequence DNA

Random-sequence DNA is the optimal form of biologically irrelevant DNA, both for measuring transcriptional noise and for identifying sequences important for a biological function in an unbiased manner. We computationally generated an 18 kb stretch of random-sequence DNA. A yeast strain containing this random-sequence DNA was generated by homologous recombination *in vivo* of introduced DNAs encoding the computationally sequence (Gibson et al. 2008; Kouprina and Larionov 2008). Specifically, overlapping oligonucleotides were co-transformed with a transformation-associated recombination (TAR) vector to assemble a ∼27 kb circular chromosome, dubbed ChrXVII, capable of propagating in yeast cells (Figure S1). For unknown reasons, the resulting strain grows slightly more slowly than the parental strain (155 min vs. 140 min doubling time in synthetic medium).

### Nucleosomes form efficiently on random-sequence DNA

As chromatin is the physiological template for transcription, assessment of biological noise requires understanding the chromatin structure of the random-sequence chromosome. Poor chromatin assembly on random-sequence DNA would likely lead to artificially high levels of transcription and hence an exaggerated level of transcriptional noise. We investigated the nucleosomal pattern by performing MNase-Seq (paired end reads) on isolated mononucleosomes and a purified DNA control. Importantly, nucleosome patterns on random-sequence and genomic DNA are analyzed in the same samples.

Biological replicates are highly reproducible, and nucleosome positions in yeast genomic DNA are indistinguishable from those reported in multiple studies (Lee et al. 2007; Brogaard et al. 2012) (Figure S2). We combined the replicates to generate a total of ∼90 million single nucleosomes and ∼40 million fragments from the naked DNA controls. Of these, ∼300,000 nucleosomes (16.6 alignments and 2350 hits per bp) and 80,000 control DNA fragments (4.5 alignments and 640 hits per bp) belong to ChrXVII, which is sufficient coverage to study chromatin structure on random-sequence DNA.

As determined by the length of the paired end sequencing reads, nucleosomes on random and genomic DNAs are structurally similar (Figure S3A). In both cases, the major class of nucleosomes average 147 bp (range 143-180), the expected size (Figure S3A, B). There is also a minor class averaging 129 bp (range 120-142) that presumably arises from MNase cleavage roughly two helical turns within the nucleosome itself. These two size classes of MNase-generate fragments, and an even more minor and smaller one (range 80-119 bp), behave similarly with respect to nucleosome positioning (Figure S3C, D). With respect to nucleosome occupancy, ChrXVII produces a different MNase-Seq profile than its purified DNA control (Figure 1A, S2C). In contrast, MNase patterns of *in vivo* and purified mitochondrial DNA are indistinguishable. This indicates that nucleosome locations on ChrXVII are not random.

**Figure 1.**
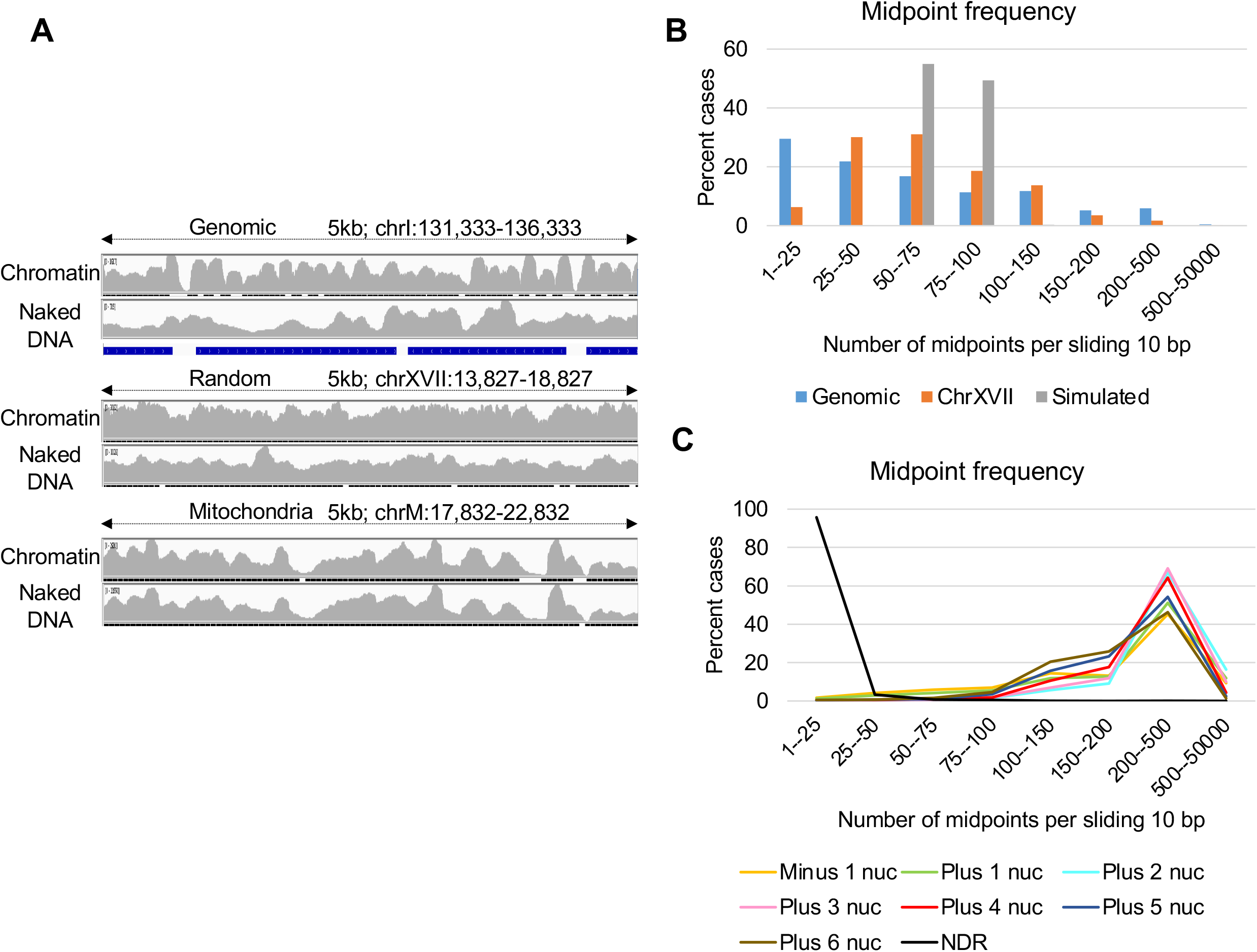
Chromatin structure on random-sequence DNA is similar, but not identical, to yeast genomic chromatin. (*A*) Browser snapshot representing 5-kb windows of aligned MNase-Seq fragments from genomic, random and mitochondrial DNA, both for chromatin and naked DNA. (*B*) Midpoint frequencies per sliding 10 bp window classified into distinct 8 categories for genomic (blue), random (ChrXVII; orange) and randomly simulated data (gray) displayed as percent of the total. (*C*) Midpoint frequency representing percent of the midpoints found within 10 bp of nucleosome depleted regions (NDRs) and the −1 to +6 nucleosome centers.

Based on the number of nucleosomal DNA fragments, it appears that there are 2.24-fold more nucleosomes on random-sequence DNA vs. genomic DNA. At first glance, this is physically impossible because the vast majority of yeast genomic DNA is associated with nucleosomes. However, there are at least three technical reasons for why this value is too high. First, based on the purified DNA sample, the ChrXVII copy number is 1.57 per haploid genome, a result in accord with a circular chromosome of this size (Bloom et al. 1983). Second, as discussed below, ChrXVII lacks nucleosome-depleted regions typical of yeast promoter and terminator regions. The sequencing read ratio for ChrXVII/intergenic is 1.80, whereas it is only 1.28 for ChrXVII/genic. Third, random-sequence DNA does not have long AT-rich stretches, and its 50% GC content is higher than the 40% of yeast genomic DNA. As MNase cleaves dinucleotides with A or T residues, nucleosomes with AT-rich sequences are preferentially degraded and hence lost from the sample, particularly when MNase cleavage generates mononucleosomes as the major product (Fan et al. 2010). Thus, these results indicate nucleosomes form on random-sequence DNA with an efficiency comparable to genomic DNA.

### Random-sequence DNA has fewer nucleosome-depleted regions and a lower degree of nucleosome positioning and phasing

To investigate nucleosome positioning, we analyzed the frequencies of nucleosome midpoints that are defined by the central position of an MNase-generated sequence. Such nucleosome midpoints depend on the precise location of MNase cleavage within flanking linker sequences, so a nucleosome located at a precise position will inevitably be observed as multiple adjacent midpoints. We therefore consolidated midpoints within a moving average of 10-bp windows. With 90 million fragments per 12 million bp yeast genome and 10-bp sliding windows, 75 nucleosome midpoints are expected to occur by chance, which is seen from the peak of the computationally simulated nucleosomes (Figure 1B).

The occurrence of midpoints per nucleotide position for genomic and random-sequence chromatin was compared to these computationally randomized midpoint locations (Figure 1B). Virtually all computationally simulated nucleosomes have 50-100 midpoints, which corresponds to random positioning. As expected from the clear pattern of nucleosome positioning in virtually all eukaryotic species, only 25% of nucleosomes in yeast genomic chromatin have 50-100 midpoints. In random-sequence chromatin, about half of the nucleosomes are poorly positioned (50% have 50-100 midpoints), and the other half shows varying levels of positioning.

Yeast chromatin is characterized by nucleosome-depleted promoter and terminator regions as well as phased and positioned nucleosomes in coding regions. As expected, 30% of nucleotide positions in the yeast genome have fewer than 25 midpoints and hence are nucleosome-depleted, whereas 6.4% have > 200 midpoints indicative of well-positioned nucleosomes (Figure 1B). The highest number of midpoints per position is associated with the +1 to +3 nucleosomes, and this value decreases gradually for nucleosomes at more downstream positions (Figure 1C).

In contrast, within random-sequence DNA, far fewer (6%) positions have < 25 midpoints, indicating a paucity of nucleosome-depleted regions (Figure 1B). Furthermore, there are considerably fewer well-positioned nucleosomes (only 1.7% have > 200 midpoints) on the random-sequence DNA than on genomic DNA. Thus, the striking chromatin pattern in yeast genomic DNA is considerably muted in random-sequence DNA. However, nucleosome positioning on the random-sequence DNA differs from that of computationally randomized positions (Figure 1B), presumably because of intrinsic DNA sequence preferences and/or spacing constraints imposed by nucleosome remodeling complexes.

To address nucleosome phasing on genomic and random-sequence DNA, we aligned the most highly positioned nucleosomes by their midpoints and analyzed midpoint frequency of adjacent nucleosomes. As expected, we observe the classic nucleosome phasing pattern on genomic DNA: i.e., multiple positioned nucleosomes with decreasing levels of positioning at increasing distance from the most positioned nucleosome (Figure 2A). In contrast, on random-sequence DNA, adjacent nucleosomes are very poorly positioned, and indeed midpoint frequencies are essentially flat after the adjacent nucleosomes (Figure 2B). In a similar vein, when regions of low nucleosome occupancy (< 25 midpoints) are aligned, regions corresponding to adjacent nucleosomes have very limited positioning. Thus, when compared to yeast genomic chromatin, ChrXVII chromatin is characterized by fewer nucleosome-depleted regions, fewer highly positioned nucleosomes, and little nucleosome phasing, although nucleosome positioning is non-random.

**Figure 2.**
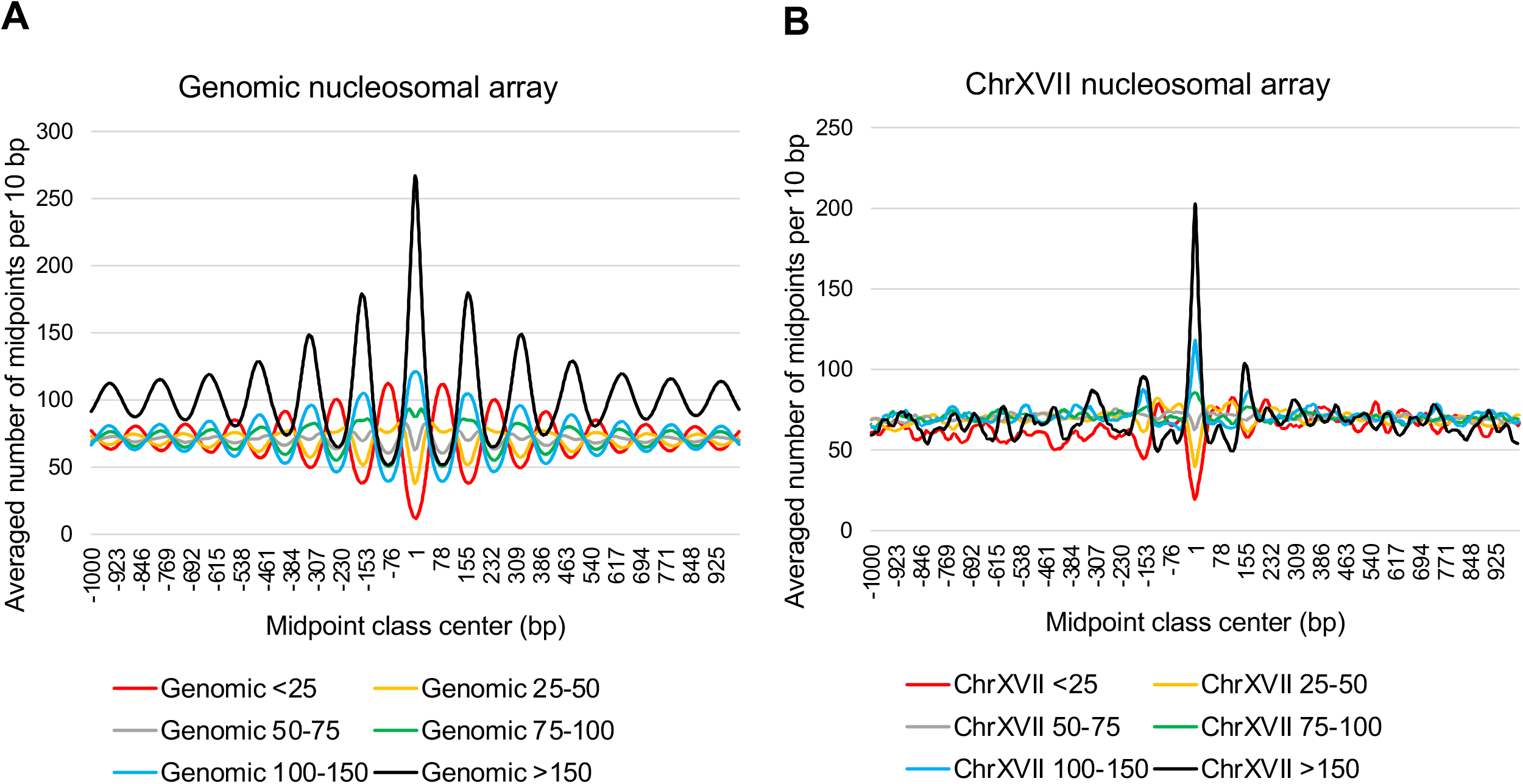
Phased nucleosome arrays form poorly on random-sequence chromatin. Meta-nucleosome position profiles of (*A*) genomic and (*B*) random-sequence DNA (ChrXVII). The average number of midpoints per 10 bp that are located 1 kb upstream or downstream of aligned nucleosomes (midpoints defined at coordinate 0) with the indicated number of midpoints (various colors).

### Levels of RNAs expressed from random-sequence DNA are comparable to those of typical yeast RNAs

We performed RNA-Seq to measure steady-state levels of poly(A)-containing RNA in the strain harboring ChrXVII. Transcription occurs readily in both directions throughout the random-sequence DNA with few, if any, regions where RNAs are absent (Figure 3A). In contrast, and as expected, mRNAs transcribed from genomic DNA are organized into distinct regions with directionality (sense gene alignments are blue and antisense are red). In addition, very little stable RNA is observed between mRNA coding regions.

**Figure 3.**
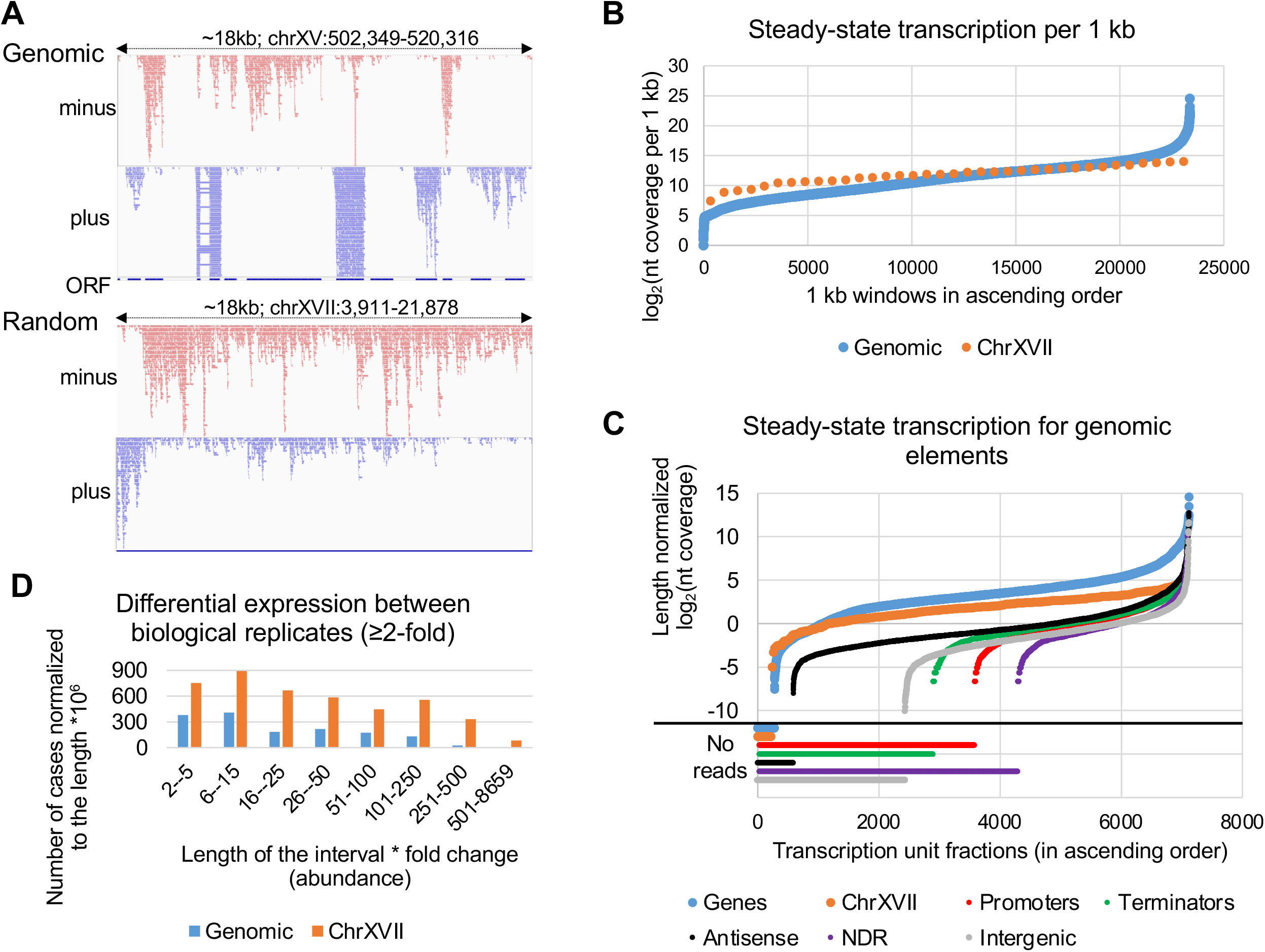
Steady-state levels of random-sequence RNAs are comparable to most yeast mRNAs but show more cell-to-cell variability. (*A*) Snapshot from the IGV browser showing aligned RNA-Seq fragments from genomic and random-sequence DNA, with sense (blue) and antisense (red) transcripts within an 18-kb window. (*B*) Steady-state levels of RNA expressed from genomic (blue) and random-sequence (orange) DNA as measured by nt coverage per 1-kb windows sorted in ascending order. (*C*) Steady-state RNA levels for ChrXVII (calculated for 100 nt windows) and for indicated (various colors) genomic elements (normalized to length). Nucleotide coverages were normalized to the lengths of all windows before deriving log_2_ values which are plotted in ascending order. (*D*) Number of cases with differential (> 2-fold) nt coverage between replicates in the indicated 8 categories defined by the length of the interval multiplied by the fold increase and normalized to the length of genomic and random-sequence DNA.

We quantified RNA levels expressed from random-sequence and genomic DNA in 1-kb windows and plotted the results as log_2_(nucleotide coverage) in ascending order (Figure 3B). As expected, most RNAs from most genomic DNA are expressed at comparable levels, although 3.6% of the genome is expressed at considerably lower levels (presumably intergenic or silenced regions) and 7.6% of the genome is expressed at considerably higher levels (primarily ribosomal and glycolytic genes). Transcription from all regions of random-sequence DNA leads to similar RNA levels that are roughly comparable to that of the majority of mRNA expressed from yeast genomic DNA (Figure 3B). When compared to various functional genomic regions, the amount of transcription at random DNA is slightly lower than that of mRNA coding regions (genes), but it is much higher than that of promoters, terminators, antisense within genes, nucleosome-depleted regions, and intergenic regions (Figure 3C). Thus, as assayed by steady state RNA levels, transcriptional noise from random-sequence DNA occurs at roughly comparable levels to those of typical mRNAs, but it is strongly suppressed at other many non-coding regions of the yeast genome.

### Transcription from random-sequence DNA is more variable than from genomic DNA

We analyzed biological replicates to estimate the extent of transcriptional variability within a population of cells for genomic and random DNA. Interestingly, the correlation coefficient for genomic-derived RNAs (0.99) is higher than for random-sequence derived RNAs (0.94). As these correlations involve the identical samples, and RNAs are expressed at roughly comparable levels, this observation suggests that transcription from random-sequence DNA is more variable. To expand upon this observation, we scanned the two replicates for intervals with at least 2-fold difference in nucleotide coverage. The length of the interval with differential coverage for the replicates depends on the continuity of ≥ 2-fold difference between adjacent nucleotides. The variation between the replicates is expressed as the fold-change multiplied by the length of the interval (Figure 3D). From this analysis, the sum of the differential intervals is 2.8 times higher for ChrXVII-encoded RNAs as opposed to genomic-encoded RNAs. Thus, transcription from random-sequence DNA is more variable than from genomic DNA.

### Transcription levels are higher and RNAs are less stable when expressed from random-sequence DNA

Measurements of steady-state RNA levels reflect a balance between synthesis and degradation. We measured transcription rates using 4tU-seq, an approach involving a 5-minute pulse of 4-thiouracil (4tU), resulting in metabolic incorporation into newly synthesized RNA (Miller et al. 2011; Sun et al. 2013; Barrass et al. 2015) that is subsequently enriched via activated disulfides conjugated to biotin (Duffy et al. 2015). Levels of newly synthesized RNA and hence rates of transcription are roughly comparable throughout the random-sequence DNA, but they are 3-fold higher than that of typical yeast mRNAs (Figures 4A and S4). Interestingly, active transcription from random-sequence DNA is much higher than from antisense, nucleosome-depleted regions, promoters, terminators, and intergenic regions (Figure 4B), suggesting that transcriptional noise from these regions is suppressed.

**Figure 4.**
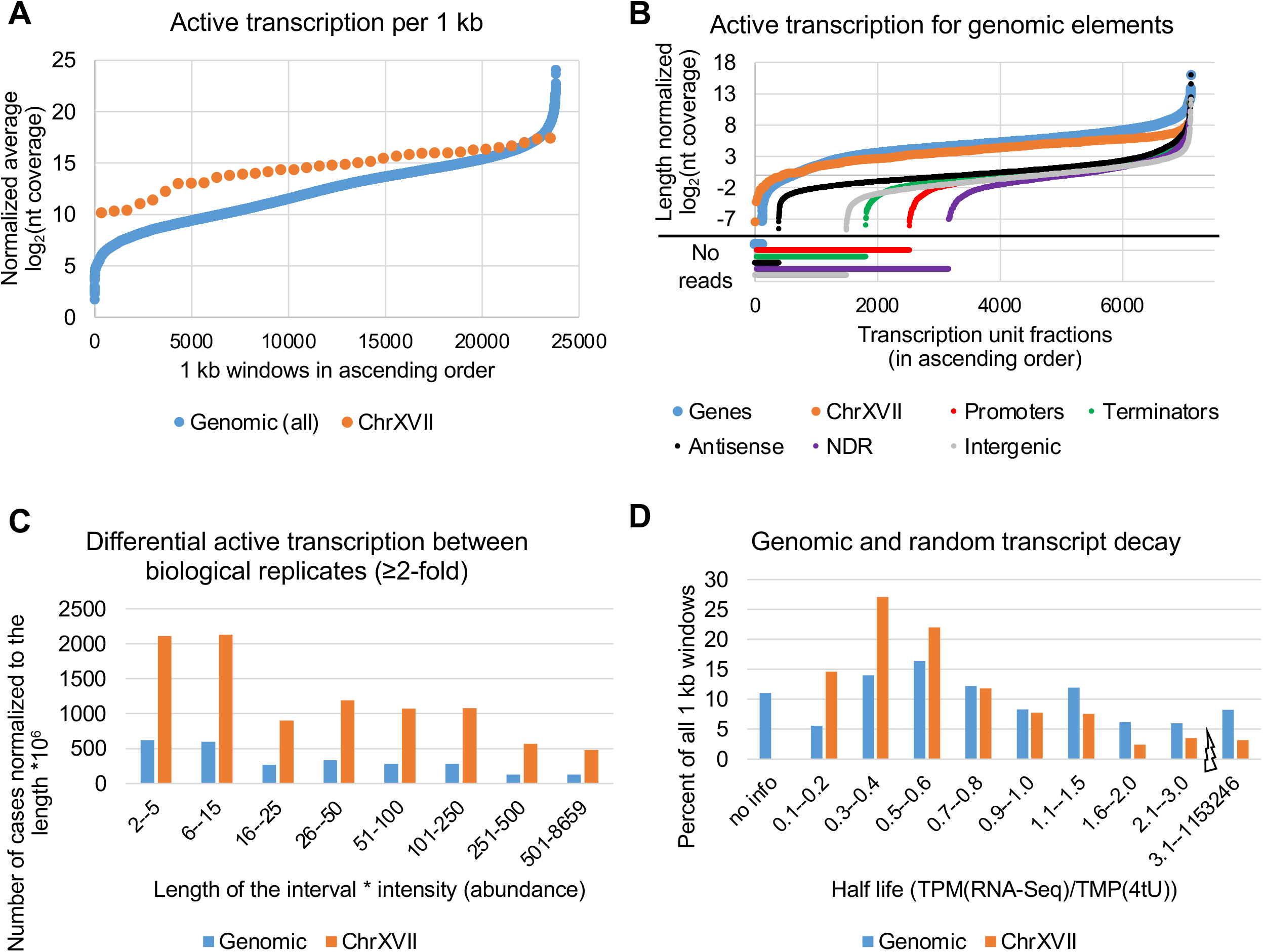
Active transcription and RNA degradation rates are higher for RNAs expressed from random-sequence vs. genomic DNA. (*A*) The amount of active transcription from genomic (blue) and random-sequence DNA (orange) as measured by background-subtracted 4tU-Seq nt coverage per 1-kb windows that are plotted in ascending order. (*B*) Levels of active transcription for the indicated genomic elements (normalized to their lengths with ChrXVII split into 100 nt windows). (*C*) Number of cases with differential (> 2-fold) nt coverage between replicates in the indicated 8 categories defined by the length of the interval multiplied by the fold increase and normalized to the length of genomic and random-sequence DNA. (*D*) RNA half-lives of genomic and random-sequence RNAs were calculated as TPM(RNA-Seq)/TMP(4tU)). The number of cases of the indicated 9 categories are shown as the percent of the total. 1 kb windows for which either RNA-Seq or 4tU-Seq = 0 were excluded (no info).

As observed with steady-state RNA levels, newly synthesized RNA and hence transcription rates within cell populations are more variable on random-sequence DNA than on genomic DNA. On newly synthesized RNA, the nucleotide-level correlation between biological replicates is 0.97 on yeast genomic DNA and only 0.86 on random-sequence DNA. In addition, the sum of differential intervals on random-sequence DNA is 3.6-fold higher than on genomic DNA (Figure 4C). A comparison of populational differences between random-sequence and genomic transcription reveals that actively transcribed RNA levels are more variable than steady-state RNA levels (Figures 3D, 4C).

Steady-state RNA levels (M) are determined by the rate of transcription (β) and the rate of RNA decay (α) according to the formula M = β/α (Schwanhausser et al. 2011; Sun et al. 2013; Blumberg et al. 2021). As the steady-state levels of RNAs generated from random-sequence DNA are roughly comparable to those of the majority of genomic-encoded RNAs (Figure 3B), this means that decay rates of RNAs expressed from random-sequence DNA are ∼3-fold faster (Figure 4D). Thus, the level of transcriptional noise is somewhat higher than the level of transcription of typical yeast genes.

### Similar, but not identical, polyadenylation patterns of RNAs expressed from random-sequence and genomic DNA

We used 3’READS (Jin et al. 2015) to compare the polyadenylation patterns of RNAs expressed from random-sequence and yeast genomic DNA (Figure 5A). As expected, polyadenylation of yeast mRNAs is almost completely restricted to 3’UTRs, with each gene having an average of ∼50 3’ isoforms (Moqtaderi et al. 2013). In contrast, RNAs expressed from ChrXVII are polyadenylated throughout the entire region of random-sequence DNA (Figures 5A and S5A, B).

**Figure 5.**
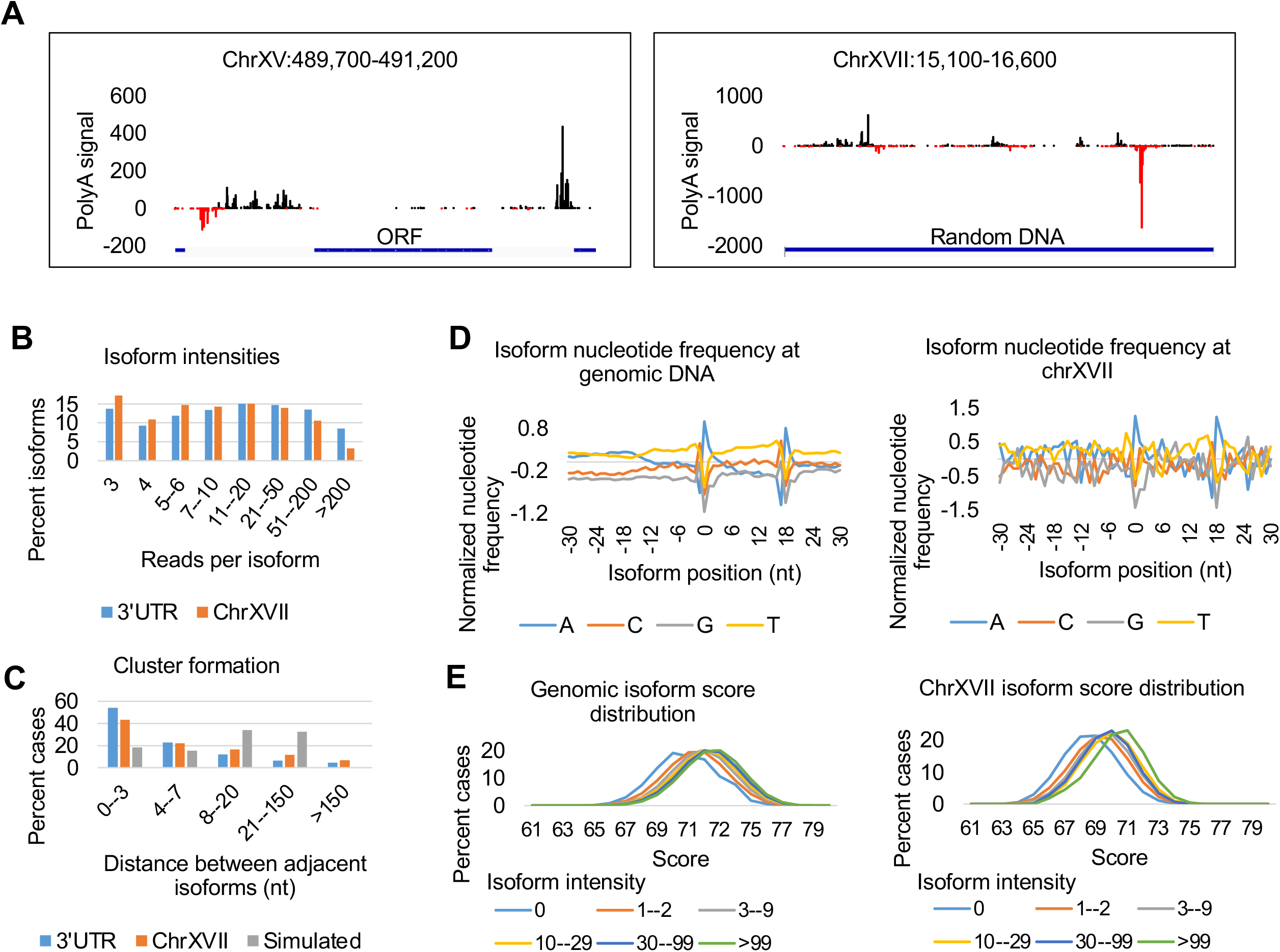
Poly(A) site profiles of random-sequence and yeast mRNAs are roughly similar. (*A*) Poly(A) reads aligned to representative 1.5 regions of genomic and random-sequence DNA. (*B*) Isoform levels (reads per isoform) for poly(A) sites in genomic 3’UTRs (blue) and random-sequence RNAs (orange). (*C*) Frequency of distances between adjacent 3’ isoforms for within 3’UTRs (blue), random-sequence RNAs (orange), and computationally simulated data that randomizes positions (gray). (*D*) Nucleotide frequencies at positions −30 to +30 for the most highly expressed (top 10%) 3’ isoforms from genomic (normalized to nucleotide frequency in the yeast genome) and ChrXVII RNAs. (*E*) Isoform score distributions (Moqtaderi et al., 2013) of genomic and ChrXVII RNAs in 6 bins (different colors) based on isoform read intensity.

To compare 3’ isoform features between RNAs expressed from random-sequence and genomic DNA, we examined the number of reads per isoform (relative isoform levels) and adjacent isoform distances (isoform clusters as compared to random simulation of isoform locations). Both the range of 3’ isoform levels (Figure 5B) and cluster formation (Figure 5C) are similar for 3’UTRs of yeast mRNAs and random-sequence RNAs. However, the maximally expressed isoform and maximum isoform intensities within 400 nt windows tend to be slightly lower for RNAs expressed from random-sequence DNA than from genomic DNA (Figure S5C, D). Consistent with these observations, the number of 3’ isoforms per 400 nt window is higher for ChrXVII RNAs than for 3’ UTRs (Figure S5E). The distance between maximum isoforms in adjacent, non-overlapping 200 nt windows is larger in RNAs from ChrXVII are larger than from genomic DNA (Figure S5F). The distance between the first and last isoforms within a 400 nt sliding window have a propensity to be below 100 nt for 3’UTRs and above 200 nt for random-sequence DNA (Figure S5G). Taken together, ChrXVII and yeast mRNAs have roughly similar polyadenylation patterns, although ChrXVII 3’ isoforms tend to be more spread out, slightly more frequent, and relatively less abundant than 3’ isoforms in yeast mRNAs.

The nucleotide composition around poly(A) sites of random-sequence RNAs mimic those within the 3’UTRs of yeast mRNAs (Figure 5D): There is a spike of A +1 to +3 nt with respect to the isoform position (63% nucleotide frequency for genomic and 80% for random); +18 to +20 nt downstream A peak for genomic (61%) and for random (42%); both +8 to +16 and −4 to −2 regions have tendency to be enriched in A residues (38-34% for genomic and 44-33% for random); comparable GC nt “rise and fall” profiles for the first −2 to +1 nt peak and +16 to +18 downstream (Figure 5D). Lastly, when compared to the nucleotide sequence matrix of yeast 3’ isoforms (Moqtaderi et al. 2013), the most highly expressed ChrXVII isoforms have scores that are nearly as high as those of yeast 3’ isoforms, and the isoform score distributions drop similarly in both random-sequence and genomic RNAs in accord with their level of expression (Figure 5E). Thus, the choice of poly(A) sites by the cleavage/polyadenylation machinery is roughly similar for RNAs expressed from random-sequence and genomic DNAs.

### 5’ isoform profiles from random-sequence DNA show very low specificity, comparable to non-5’UTR regions of the yeast genome

We mapped 5’ isoforms by modifying a previously described dephosphorylation-decapping method (Machida and Lin 2014; Pelechano et al. 2016) and obtained the expected metagene profile for endogenous yeast genes (Figure S6A). 5’ isoforms emanating from ChrXVII are scattered at low levels throughout the entire stretch of 18 kb of random-sequence DNA with few, if any, well expressed isoforms (Figure 6A). This pattern is very different from 5’ isoforms from genomic DNA, which heavily biased to promoter regions (Figure 6A, left). However, increased magnification reveals that random-sequence DNA and genomic regions outside 5’UTRs express many 5’ isoforms at low levels (Figure 6A, right). We then measured the number of 5’ isoforms per various functional genomic elements, including the whole genome split into 200 nt windows. The 5’ isoform pattern from random-sequence DNA resembles that observed from yeast genomic regions not corresponding to 5’UTRs, and it is slightly lower than within ORFs (Figures 6A, B and S6B). This suggests that transcriptional noise is initiated with very low specificity and that 5’ isoforms initiated from non-5’UTRs within genomic DNA are likely to be transcriptional noise. TSS nucleotide frequencies for ChrXVII 5’ isoforms display some similarities to 5’ isoforms of yeast mRNAs (Figure S6B). Unlike the case for 3’ isoforms, which are typically clustered, the distances between adjacent 5’ isoforms for ChrXVII RNAs, are largely expected by chance, indicating weak clustering (Figure S6D).

**Figure 6.**
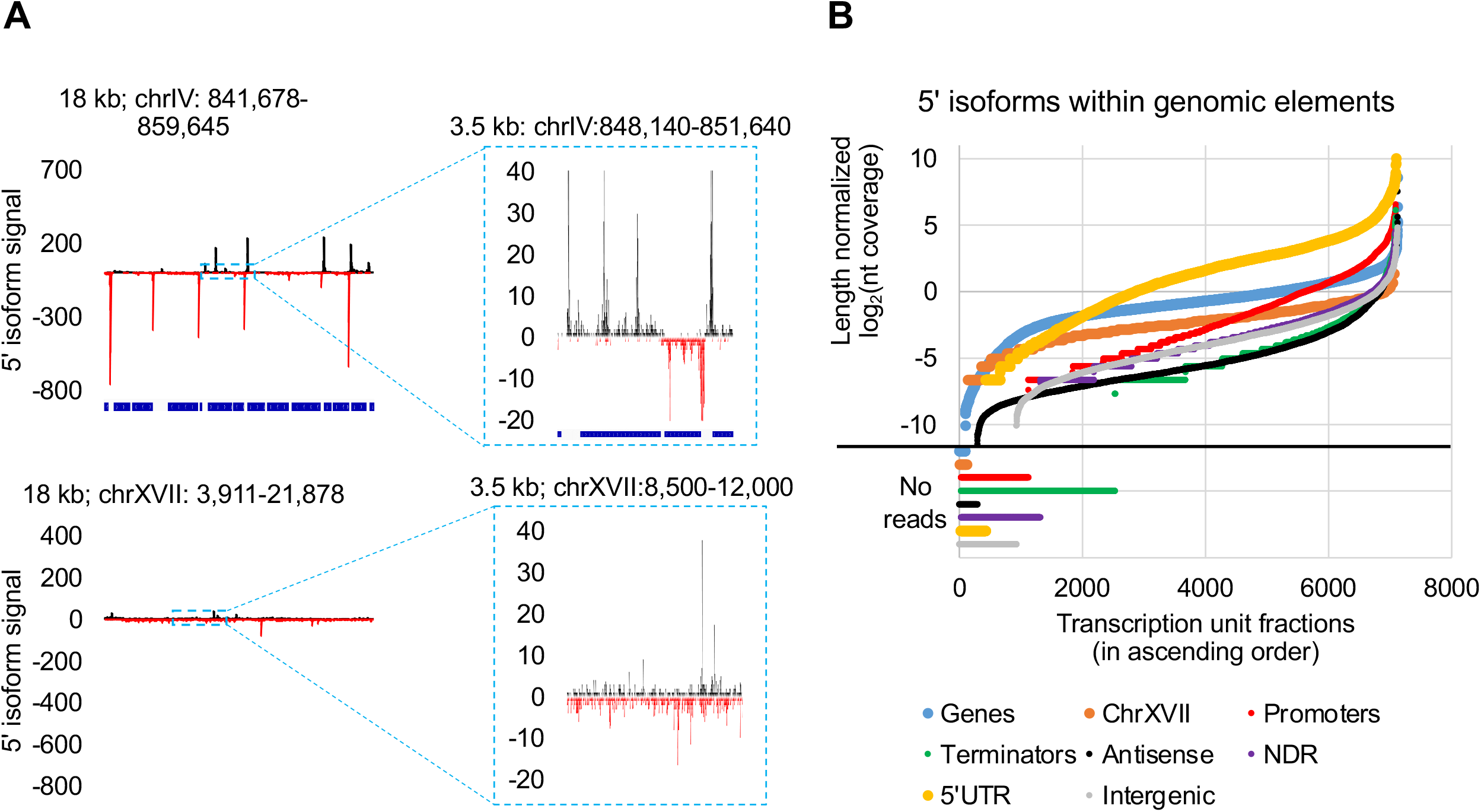
Numerous 5’ isoforms expressed from random-sequence DNA and from yeast genomic regions outside 5’ UTRs. (*A*) Aligned 5’ isoform reads for genomic and random-sequence RNA within 18 kb (left) and 3.5 kb (right; enlarged by a factor of 20) windows. (*B*) Number of 5’ isoforms for random-sequence RNAs (calculated for 100 nt windows) and for various functional genomic regions. Nucleotide coverages were normalized to the lengths of all windows before deriving log_2_ values and plotting the values in ascending order.

## DISCUSSION

### Studying biological noise using random-sequence DNA

Previously, we addressed the issue of biological function by assaying nucleosome positioning (Hughes et al. 2012), transcription (Jin et al. 2017), and polyadenylation (Moqtaderi et al. 2013) in the yeast *S. cerevisiae* on evolutionarily irrelevant DNA from the distant yeast species *D. hansenii*. However, while *D. hansenii* DNA is evolutionarily irrelevant in *S. cerevisiae* cells, it is not evolutionarily neutral, and hence not suitable for measuring biological noise. Here, we measure biological noise in yeast cells by analyzing chromatin structure and various aspects of transcription of an 18 kb region of DNA whose sequence was randomly generated. Molecular events on random-sequence DNA are not only functionally irrelevant *per se*, but we assume that they mimic biological noise that occurs on yeast genomic DNA. Conversely, we presume that events that occur on genomic, but not random-sequence, DNA are biologically meaningful. In addition, this approach complements, but is distinct from, experiments that define sequences required for a specific molecular function via genetic or biochemical selections from libraries of molecules with short stretches of random DNA sequences.

### Chromatin structure on random-sequence DNA

Essentially all of ChrXVII is nucleosomal, but chromatin on random-sequence DNA does differ from chromatin of yeast genomic DNA. First, genomic DNA has many more regions of low nucleosome occupancy than random-sequence DNA, providing direct evidence that such regions are subject to evolutionary selection and hence functional. Indeed, these nucleosome depleted regions are almost always in promoters and terminators. Second, chromatin on random-sequence DNA shows a lower degree of nucleosome positioning, but there are some highly positioned nucleosomes, presumably due to DNA sequence specificity for nucleosome formation (Drew and Travers 1985). Third, nucleosomes on random-sequence DNA are poorly phased, even when flanking a highly positioned nucleosome, arguing against a simple statistical positioning model (Fedor et al. 1988).

Nucleosome-depleted regions of the yeast genome arise from 1) poly(dA:dT) sequences (Struhl 1985), which have intrinsic nucleosome destabilizing properties (Segal and Widom 2009; Zhang et al. 2009; Hughes et al. 2012) and are preferred substrates of the RSC nucleosome remodeling complex (Lorch et al. 2014; Krietenstein et al. 2016), and 2) binding sites for activator proteins that recruit nucleosome modelers and histone acetylase complexes. Nucleosome-depleted regions support localized transcriptional initiation that leads to a highly positioned +1 nucleosome followed by phased but progressively less positioned nucleosomes further downstream (Struhl and Segal 2013). This nucleosome positioning pattern and bidirectional transcription occurs even when nucleosome-depleted regions are generated fortuitously on evolutionarily irrelevant DNA (Hughes et al. 2012; Jin et al. 2017). Importantly, the specific locations of functional sequences within the genome results in a nucleosome positioning pattern that is similar (though not identical) among cells in a population.

In contrast, random-sequence DNA lacks functional elements at specific locations, and most nucleosomes are weakly positioned. As such, nucleosome positions on ChrXVII are very likely to differ considerably from cell to cell, except for the small minority of well-positioned nucleosomes. Furthermore, and as discussed below, transcription is initiated at numerous sites throughout ChrXVII, yielding RNAs that differ considerably among cells in the population. Thus, while the classic nucleosome positioning pattern should occur upon transcription from a given site in an individual cell, the overall cell population will have +1 and subsequent nucleosomes at numerous positions. In addition, highly positioned nucleosomes on ChrXVII reflect intrinsic sequence preferences of histone octamers, which is insufficient for nucleosome phasing. Formation of phased nucleosome arrays requires transcriptional initiation from a defined location (Struhl and Segal 2013) and transcription-dependent spacing constraints of the Chd1 and Isw1 nucleosome remodelers (Gkikopoulos et al. 2011).

### Transcriptional noise in yeast occurs at high levels but is suppressed at some genomic locations

The nucleosomal nature of chromosome XVII is important for interpreting the transcriptional properties of random-sequence DNA and hence measurements of transcriptional noise. As expected for random-sequence DNA, steady state RNA levels are roughly similar across the 18 kb of ChrXVII. Interestingly, the steady-state levels of RNAs transcribed from random-sequence DNA are comparable to those of most yeast mRNAs. Moreover, levels of newly synthesized random-sequence RNAs (4tU-seq) are ∼4-fold higher than those of typical mRNAs, indicating a high level of transcriptional noise. The difference between steady state and newly synthesized RNA levels indicates that RNAs generated from random-sequence DNA are considerably less stable than conventional mRNAs.

The relative instability of random-sequence RNAs is likely due to at least two reasons. First, 3 out of 64 codons are nonsense codons, so translation of random-sequence RNAs will generally yield very short proteins, hence subjecting these RNAs to nonsense-mediated decay (He and Jacobson 2015; Kurosaki et al. 2019). Second, due to the first ATG rule for translational initiation and the fact that only 1 of 64 codons are bound by the methionine initiator tRNA, many 5’ UTRs are likely to be long and possibly subject to increased degradation (Chan et al. 2018).

The level of newly synthesized RNA from random-sequence DNA is in excellent accord with calculated estimates that only 10-20% of elongating Pol II molecules generate mRNAs, such the remaining 80-90% represent transcriptional noise (Struhl 2007). However, unlike random-sequence RNAs that are synthesized across ChrXVII at roughly comparable levels to those of yeast mRNAs, synthesis of antisense RNAs and transcripts from nucleosome-depleted regions, promoters, terminators, and intergenic regions occurs at much lower levels. In addition, many individual regions of the yeast genome show little if any transcription. These observations strongly suggest that yeast cells have a mechanism(s) to suppress biological noise at many genomic locations, including those (e.g., promoters and terminators) that have biological function in the native organism.

We are unable to distinguish between synthesis of true mRNAs and biological noise on the sense strand within mRNA coding regions. It is likely that some of the measured level of synthesis within mRNA coding regions is due to internal initiation. This suggestion is consistent with our observation of many 5’ ends within coding regions and the existence of many internal transcripts that are only observed in strains defective in mRNA degradation (Arigo et al. 2006; Thiebaut et al. 2006; Rougemaille and Libri 2010). In this regard, we speculate that chromatin structural changes associated with transcription of classically defined mRNAs permits increased levels of internal transcription.

### Inherent specificity of transcriptional initiation and polyadenylation

In yeast, 5’ and 3’ ends are generated by independent molecular processes that are not mechanistically connected (Pelechano et al. 2013). Profiles of 5’ and 3’ ends of random-sequence RNAs, respectively, provide information on the inherent specificity of transcriptional initiation and polyadenylation that is unbiased by evolutionary selection. Although 5’ ends of RNAs are not randomly distributed across ChrXVII, there are numerous 5’ ends and no strongly favored initiation sites. These results indicate that the inherent specificity of transcriptional initiation by the basic Pol II machinery is low. For polyadenylation, only a small subset of potential poly(A) sites is used, and there is a considerable range in the levels of 3’ isoforms. This indicates that the inherent specificity of polyadenylation is higher than that of transcriptional initiation, although the large number of distinct poly(A) sites suggests a modest level of specificity.

### Evolutionary and mechanistic implications for transcription in yeast cells

Unlike the case for random-sequence DNA, transcriptional initiation and poly(A) sites are, respectively, strongly biased to promoters and 3’UTRs. For transcriptional initiation, this bias reflects poly(dA:dT) sequences and activator binding sites that lead to nucleosome-depleted regions that create windows for the Pol II machinery (Hughes et al. 2012; Jin et al. 2017). Within these open windows, DNA sequences make only a minor contribution to where transcription is initiation. This limited specificity is consistent with previous observations in yeast that 1) TATA element quality is relatively unimportant for low and moderate levels of constitutive transcription typical of yeast genes (Struhl 1986; Iyer and Struhl 1995a), and 2) the Initiator element makes only a modest contribution to the level of transcription, although it is important for 5’ site choice (Chen and Struhl 1985; Hahn et al. 1985; Nagawa and Fink 1985). An important exception to this limited specificity is the importance of a consensus TATA element for high levels of transcription mediated by activator proteins under appropriate environmental conditions (Iyer and Struhl 1995a).

For polyadenylation, the mechanism of strong 3’ bias for polyadenylation is poorly understood, but it is highly likely to reflect evolutionarily selected sequences in the 3’ UTR. In yeast, the most likely candidate is the long AT-rich stretch that immediately follows the open reading frame (Lui et al. 2022). In many other organisms, an AAUAAA motif plays a key role in poly(A) site selection (Proudfoot and Brownlee 1976; Wickens and Stephenson 1984; Proudfoot 2011), but such motifs occur throughout the genome, and hence can’t be the sole basis of 3’ UTR bias and poly(A) site selection. The overall similarity in polyadenylation profiles of random-sequence RNAs and yeast mRNAs suggests that the choice of poly(A) sites primarily reflects intrinsic sequence preferences of the cleavage/polyadenylation machinery, not evolutionarily selected sequences. However, there is some evolutionary selection for poly(A) sites because poly(A) sites in random-sequence RNAs tend to be more spread out, slightly more frequent, and relatively less abundant than 3’ isoforms in yeast mRNAs.

Cleavage/polyadenylation of yeast mRNAs does not occur within long protein-coding regions, and it is associated with a large structural motif that includes a long AT-rich stretch (Moqtaderi et al. 2013). As such, the frequency of poly(A) sites in RNAs transcribed from random-sequence DNA is surprising. One possibility is that phased nucleosome arrays on transcribed regions inhibit cleavage/polyadenylation. As random-sequence chromatin lacks such phased arrays, perhaps cleavage/polyadenylation occurs on chromatin with longer linker regions between nucleosomes. Alternatively, but not mutually exclusive, mRNA translation might indirectly affect the poly(A) profile by selective degradation, and in this regard random-sequence RNAs have longer 5’UTRs (due to the low frequency of ATG codons), shorter translation products (due to high frequency of stop codons), and hence longer untranslated regions. As poly(A) profiles are based on steady-state RNA levels, perhaps the observed 3’ isoforms of random-sequence RNAs represent a subset with short 5’ and 3’ UTRs.

### Functional elements constrain cell-to-cell variability

For all RNA analyses described here, correlation coefficients of biological replicates are clearly higher for yeast genomic DNA than for random-sequence DNA. This dichotomy cannot be due to a technical issue because the correlation coefficients are derived from the identical samples. Instead, this observation suggests that RNA profiles of individual cells within a population are more variable on random-sequence DNA than on yeast genomic DNA. We suggest that this reflects the fact that functional elements in promoters and terminator regions create windows for transcriptional initiation and polyadenylation that are the same in every cell. These windows represent functional constraints that limit the variability of RNA profiles among individual cells within a population.

## MATERIALS AND METHODS

### Assembly of random sequence chromosome

Random DNA sequence was generated computationally and split in 1 kb fragments, with 100 bp overlap from each end, and these sequence fragments were synthesized by Thermo Fisher Scientific. The constructs were assembled *in vivo* using transformation associated recombination (TAR) cloning (Gibson et al. 2008; Kouprina and Larionov 2008). Briefly, overlapping DNA fragments (40 ng per 1 kb fragment) and 20 ng *Eco*RI-cut TAR vector (pCC1BAC-LCyeast) were co-transformed in yeast spheroplasts from strain BY4741 (Winston et al. 1995) to generate a circular chromosome *via* homologous recombination (Kourennaia et al. 2005). The chromosome was verified with pairs of diagnostic primers that detect successful homologous recombination at the junctions (Figure S1). The final 30 kb circular chromosome was isolated from yeast and electroporated into *E. coli* (strain DH10B, Invitrogen), and re-transformed into BY4741 using the standard lithium acetate procedure.

### Nucleosome mapping by MNase-seq

Crosslinked spheroplasts from 100 ml cultures were treated with 1600 U MNase (Worthington Biochemical Corporation) for 1-1.5 h at 37°C and mononucleosomal-sized DNA was prepared following gel electrophoresis. As a control, genomic DNA was treated with MNase and mono-nucleosome-sized fragments isolated. Library preparation was performed with minor modifications of a procedure described previously (Wong et al. 2013), and samples were sequenced with Illumina NextSeq High for 75 cycles (paired end reads).

The reads were aligned to the reference gnome, sacCer3 (UCSC, 2011) containing ChrXVII (pCC1BAC-LCyeast-20-kb-random-DNA) using bowtie2 (Langmead and Salzberg 2012), and Bam files were converted to bed using bedtools bedtobam (v.2.24.0) (Quinlan and Hall 2010). These bed files were used to compute fragment locations for correctly aligned paired-end reads, fragment size distribution and size separation, fragment midpoints locations, midpoint frequency distributions, midpoint background subtractions, moving averages, MNase cutting frequencies, deriving +2 to +6 nucleosomes, meta-analysis of phased nucleosomal arrays, and nucleosomal/NDR nucleotide scores. MNase cutting frequency was calculated for genomic and random-sequence chromatin and naked DNA by counting per nt MNase-Seq fragment ends and normalizing to the total genomic or ChrXVII values.

### Measuring steady-state RNA levels by RNA-seq

Total RNAs from two biological replicates were subjected to poly(A) selection and fragmentation using Magnesium RNA Fragmentation Module (New England Biolabs). Libraries were prepared as described previously (Jin et al. 2015), and subjected to the paired-end, strand specific high-throughput sequencing with Illumina NextSeq Mid for 150 cycles (75 nt read length) equally apportioned between R1 and R2. RNA-seq reads (10.4 and 8.1 million reads for the two replicates) were mapped to the *S. cerevisiae* genome and the introduced random sequence using bowtie2 −5 25 −3 25 –no-mixed –no-discordant –dovetail, and STAR (Dobin et al. 2013) with –quantMode TranscriptomeSAM GeneCounts – outFilterType BySJout –outFilterMultimapNmax 1 –alignIntronMin 10 –alignIntronMax 2500 –alignMatesGapMax 2500 –alignSJDBoverhangMin 1. Per nucleotide coverage from the fragment locations and counts per defined intervals were then derived.

### Measuring RNA synthesis by 4tU-seq and half-life calculations

Yeast cells containing ChrXVII were treated with either 4-thiouracil (4tU, Sigma Aldrich; 5 mM final concentration) or DMSO as a control for 5 min. Spike-in control cells include untreated *S. pombe* cells and *K. lactis* cells treated with 4tU for 5 min. After combining cells from the three species, poly(A) RNA was biotinylated with activated disulfide methane thiosulfonate (MTS) biotin (Biotium Biotin-XX MTSEA). 4tU-containing RNA was enriched as previously described (Duffy et al. 2015) and used to make libraries. From five biological replicates we obtained ∼20 million paired-end sequencing reads (excluding spike-ins). Spike-in normalized nt coverages were used to subtract background from the 4tU sample to obtain active transcription measurements.

Half-life calculations were performed from the combined RNA-Seq and 4tU-Seq values (Schwalb et al., 2016, Blumberg et al., 2021). Briefly, unitless constant of RNA half-life was estimated from the number of transcripts (RNA-Seq) divided by the production of new mRNA (4tU-Seq), with RNA-Seq and 4tU-Seq expressed as transcripts per million (TPMs):

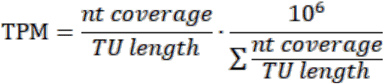

### Mapping polyadenylation sites by 3’READS

3’READS was performed as described previously (Jin et al. 2015), and we obtained ∼18 million paired-end sequence reads from 6 biological replicates. The number of isoform reads for each isoform was normalized to the transcript coverage per 50 nt window (± 25 nt from the isoform location. Using sliding windows, maximum isoforms were identified and following associated features were quantified: percent signal at the most used (maximum) coordinate; maximum isoform intensities; distances between maximum isoforms; distance between the first and the last isoform; and isoform frequencies (number of distinct coordinates). Alone standing isoforms were excluded from the sliding windows. *De novo* isoform-specific nucleotide frequencies were calculated and isoform scores were based on isoform frequency matrices (Moqtaderi et al. 2013).

### Mapping 5’ ends

Poly(A)-containing RNA (DNase I treated) was fragmented with NEBNext® Magnesium RNA Fragmentation Module, dephosphorylated with Quick CIP (NEB), heat inactivated, and ligated to a pre-adenylated 3’ adapter (Jin et al. 2015). The resulting products were treated or not treated with mRNA de-capping enzyme (New England Biolabs) and then ligated to a 5’ adapter (Jin et al. 2015), reversed transcribed, PCR amplified, and then paired-end sequenced with NextSeq Mid and NextSeq High. From 5 replicates, we obtained 32.7 million sequence reads for the de-capped sample and 25.9 million reads for the control sample. The 5’ isoforms were determined by the first position past the 5’ adapter after which we computed the 5’ reads per genomic coordinate. Negative control values per genomic coordinate were normalized using the internal *S. pombe* control and subtracted from the values of the de-capped sample.

### Data access

All raw and processed sequencing data generated in this paper have been submitted to the NCBI Gene Expression Omnibus (GEO; http://www.ncbi.nlm.nih.gov/geo/) database under accession number GSE216450.

## ACKNOWLEDGEMENTS

We thank Dan Gibson for kindly providing pCC1BAC-LC yeast vector and Joseph Geisberg and Zarmik Moqtaderi for helpful discussion. Work performed by Z.B., an employee of Google LLC, was performed during his personal time. This work was supported by a postdoctoral fellowship to Z.G. (1F32GM140555) and research grants to K.S. (GM30186 and GM131801) from the National Institutes of Health.

## AUTHOR CONTRIBUTIONS

K.S. proposed and supervised the project. Z.G. designed and conducted experimental and theoretical work and analyzed data. Z.G. and Z.B. wrote the python scripts. Z.G. and K.S. wrote the paper.

## SUPPLEMENTAL FIGURE LEGENDS

**Figure S1.**
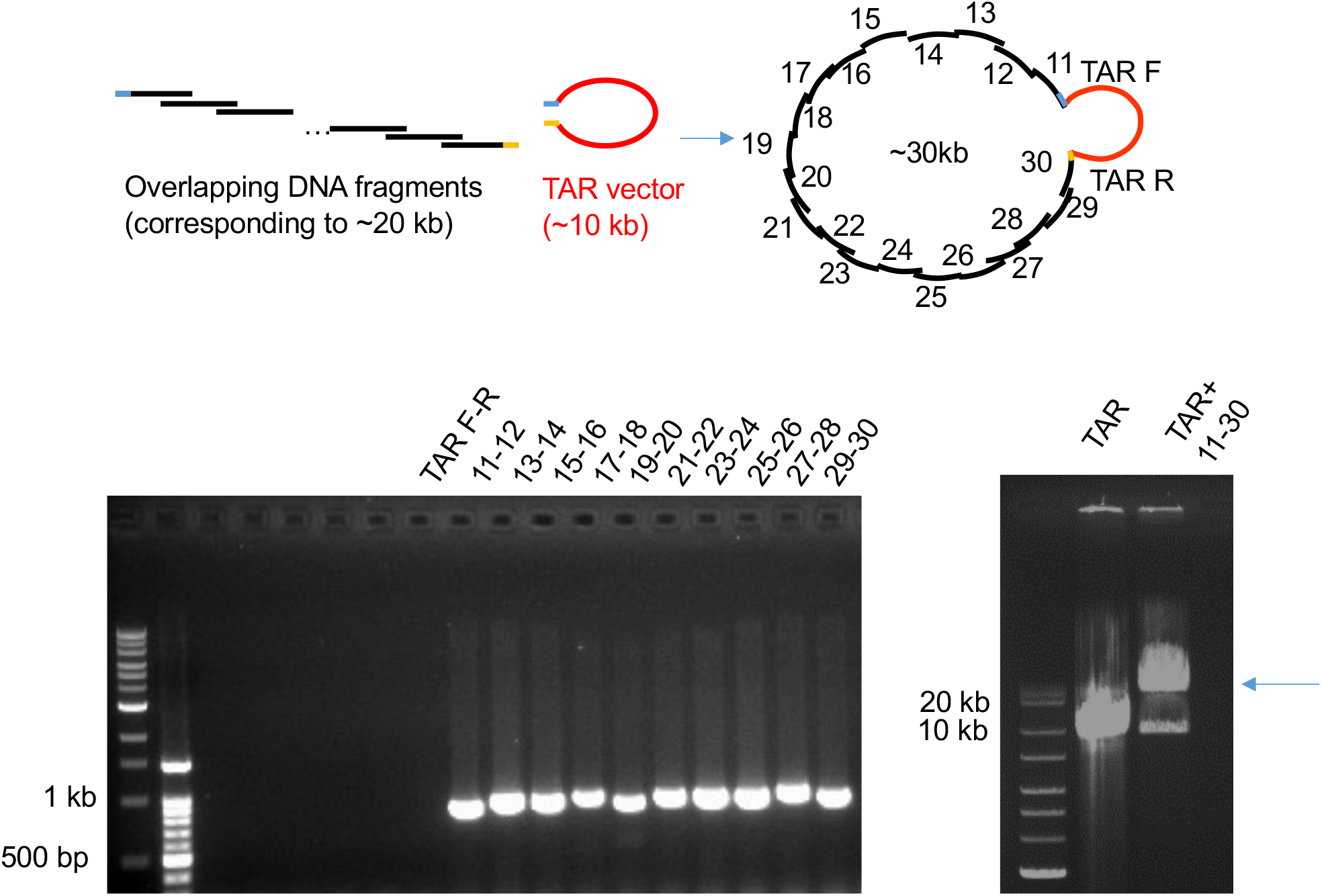
Construction of the yeast strain harboring a random-sequence chromosome. Overlapping DNA fragments, each containing a predefined, computationally randomized sequence, and the TAR vector were introduced into yeast cells with chromosome assembly occurring *in vivo* via homologous recombination (top panel). The assembled chromosome was verified with PCR amplification of the indicated fragment junctions (bottom left panel). For overall size verification, the final construct was excised from the TAR vector with the restriction sites at random-sequence ends (bottom right panel).

**Figure S2.**
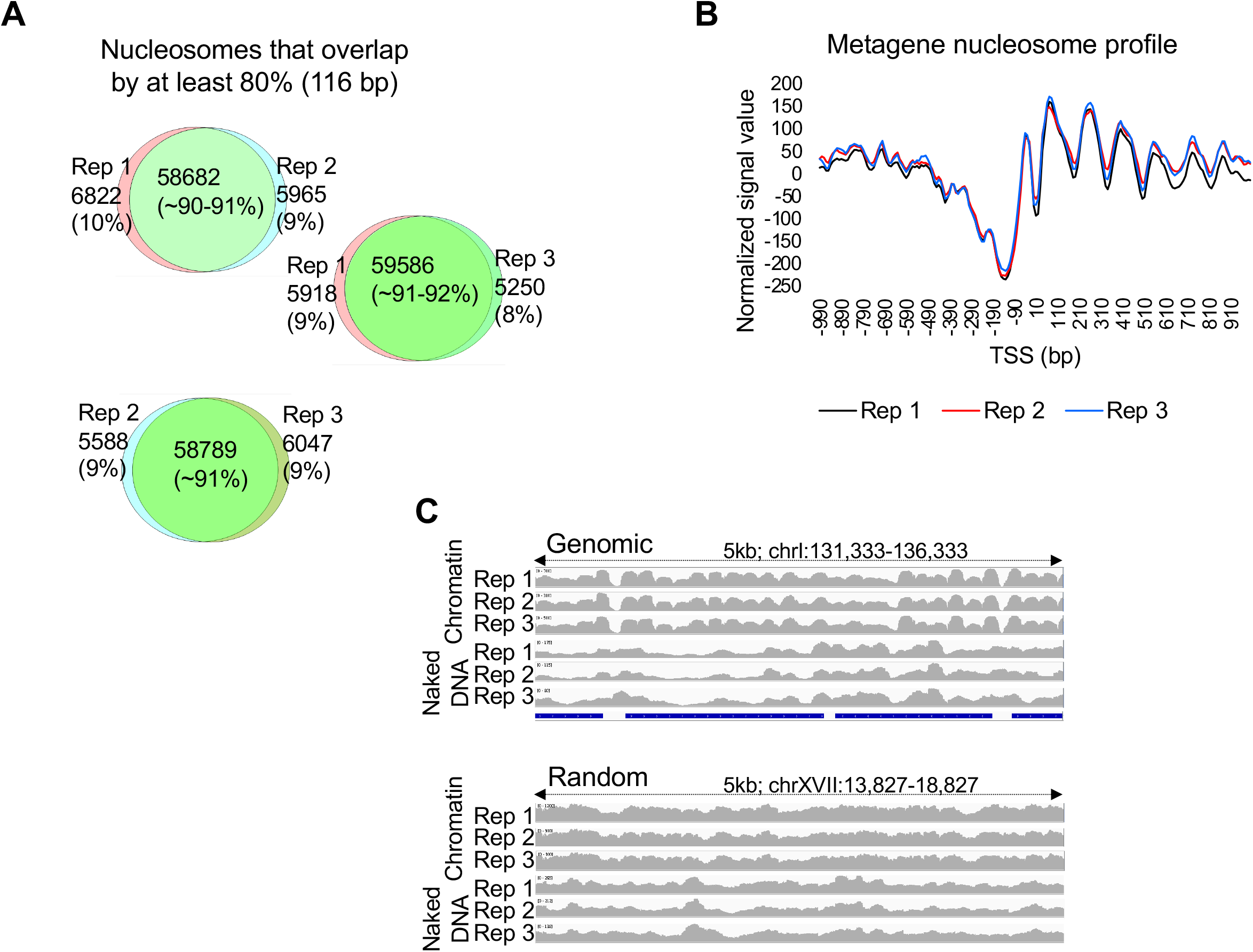
Evaluation of MNase-Seq replicates. (*A*) Overlapping nucleosome positions in pairs of biological replicates. Nucleosomes with intersecting coordinates > 116 bp (80% in coverage; 90% of all nucleosomes) between pairs of biological replicates were considered to be the same. (*B*) Nucleosome metagene profile for three biological replicates generated by aligning normalized MNase-Seq reads with respect to TSS are similar to each other and to previously reported profiles (Lee et al., 2007, Brogaard et al., 2012). (*C*) Snapshot of 5-kb windows of genomic, random-sequence, and mitochondrial chromatin and naked DNAs of 3 biological replicates.

**Figure S3.**
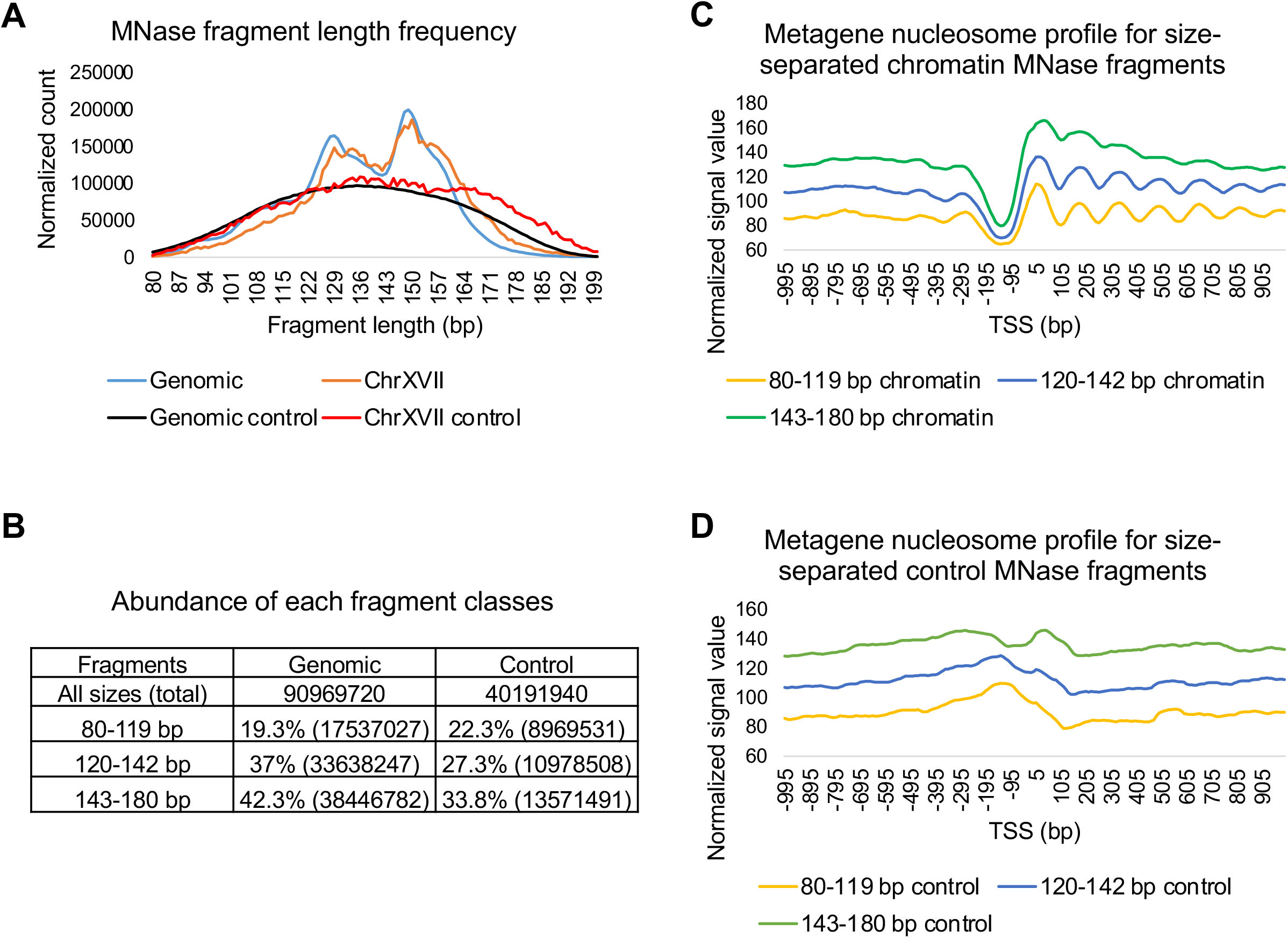
Assessment of MNase-Seq fragment classes. (*A*) Distribution of MNase-Seq fragment lengths for genomic and random-sequence (ChrXVII) chromatin and naked DNA. (B) Number of MNase-Seq reads in the indicated size classes of chromatin and naked DNA control. (*C*) Metagene nucleosome profiles of the three different MNase-seq size classes. (*D*) Metagene profiles of the indicated size classes of MNase-Seq fragments of the naked DNA controls.

**Figure S4.**
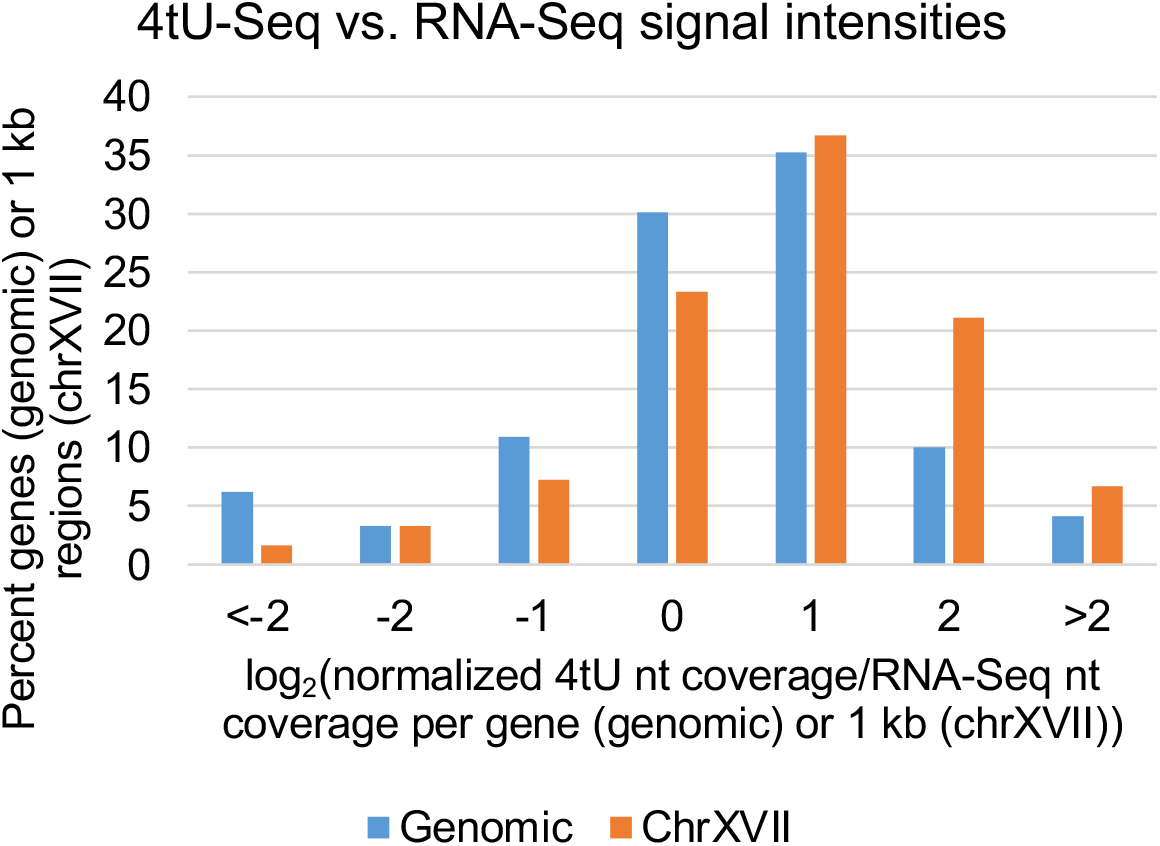
Active transcription from random-sequence DNA is higher than from genomic DNA. Percent of the genes (for genomic) or 1 kb regions (for ChrXVII) with differential active to steady-state transcript levels expressed as log_2_(4tU-Seq/RNA-Seq).

**Figure S5.**
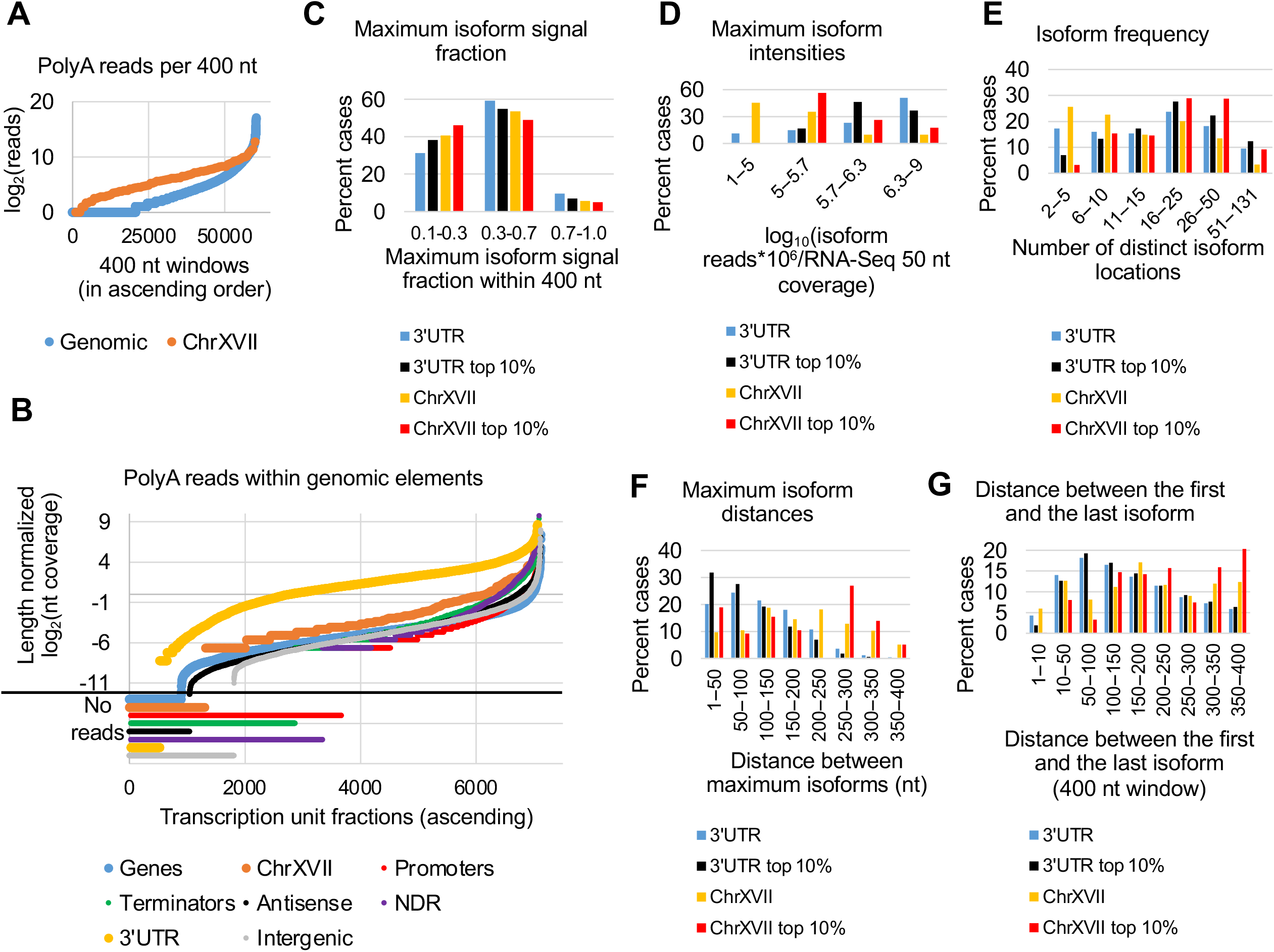
Properties of genomic and random-sequence 3’ mRNA isoforms. (*A*) Number of poly(A) reads (log_2_) per 400 nt windows in ascending order for genomic (blue) and random-sequence RNAs (normalized for the chromosome copy number; orange). (*B*) Number of poly(A) reads per 100 nt windows for the indicated functional genomic elements. (*C*) Percent of cases having the indicated ranges of maximum isoform signals (in percent of total signals) within 400 nt sliding windows in 3’UTRs and random-sequence RNAs (top 10% refers to the most highly expressed isoforms). Only isoforms with > 2 reads and isoform values after normalization to transcription were included in the analysis. (*D*) Percent of cases having the indicated ranges of maximum isoform intensities within 400 nt sliding windows in 3’UTRs and random sequence RNAs (total and top 10%). (*E*) The number of 3’ isoforms within the windows. (*F*) Distances between maximum isoforms within 3’UTR and ChrXVII within non-overlapping adjacent 200 nt windows for every nt as a starting window. (*G*) Distance between first and last isoform within 400 nt sliding windows.

**Figure S6.**
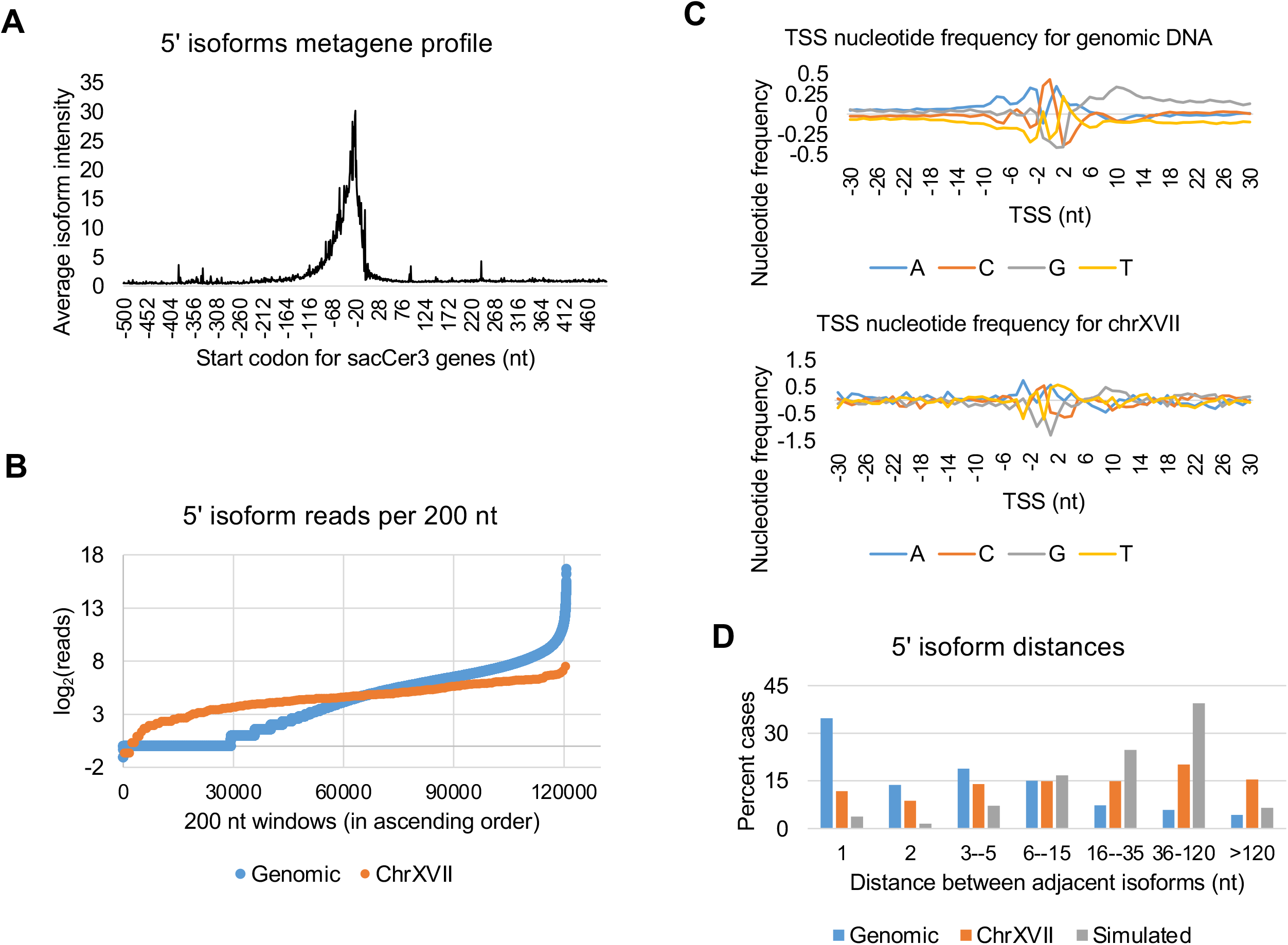
Properties of 5’ isoforms for random-sequence and genomic RNAs. (*A*) Metagene 5’ isoform signal with respect to the start codon for all yeast genes. (*B*) 5’ isoform reads per 200 nt windows for genomic (blue) and random-sequence RNAs (normalized for the chromosome copy number; orange). (*C*) Nucleotide frequencies (normalized to each nucleotide in the genome) of the region ±30 nt adjacent to the transcription start site (TSS) for 5’ isoforms expressed from genomic and random-sequence DNA with > 3 reads per isoform after background subtraction. (*D*) The frequency of distances between adjacent 5’ isoforms for genomic (blue), random-sequence (orange) and computationally randomized RNAs (gray).

## REFERENCES

Arigo JT, Eyler DE, Carroll KL, Corden JL. 2006. Termination of cryptic unstable transcripts is directed by yeast RNA-binding proteins Nrd1 and Nab3. Mol Cell 23: 841–851.

Barrass JD, Reid JE, Huang Y, Hector RD, Sanguinetti G, Beggs JD, Granneman S. 2015. Transcriptome-wide RNA processing kinetics revealed using extremely short 4tU labeling. Genome Biol 16: 282.

Bloom KS, Fitzgerald-Hayes M, Carbon J. 1983. Structural analysis and sequence organization of yeast centromeres. Cold Spring Harbor Symp. Quant. Biol. 47: 1175–1185.

Blumberg A, Zhao Y, Huang YF, Dukler N, Rice EJ, Chivu AG, Krumholz K, Danko CG, Siepel A. 2021. Characterizing RNA stability genome-wide through combined analysis of PRO-seq and RNA-seq data. BMC Biol. 19: 30.

Brogaard KR, Xi L, Wang JP, Widom J. 2012. A map of nucleosome positions in yeast at base-pair resolution. Nature 486: 496–501.

Carrozza MJ, Li B, Florens L, Suganuma T, Swanson SK, Lee KK, Shia WJ, Anderson S, Yates J, Washburn MP et al. 2005. Histone H3 methylation by Set2 directs deacetylation of coding regions by Rpd3S to suppress spurious intragenic transcription. Cell 123: 581–592.

Chan LY, Mugler CF, Heinrich S, Vallotton P, Weis K. 2018. Non-invasive measurement of mRNA decay reveals translation initiation as the major determinant of mRNA stability. Elife 7: e32536.

Chen W, Struhl K. 1985. Yeast mRNA initiation sites are determined primarily by specific sequences, not by the distance from the TATA element. EMBO J 4: 3273–3280.

Core LJ, Martins AL, Danko CG, Waters CT, Siepel A, Lis JT. 2014. Analysis of nascent RNA identifies a unified architecture of initiation regions at mammalian promoters and enhancers. Nat Genet 46: 1311–1320.

David L, Huber W, Granovskaia M, Toedling J, Palm CJ, Bofkin L, Jones T, Davis RW, Steinmetz LM. 2006. A high-resolution map of transcription in the yeast genome. Proc. Natl. Acad. Sci. U.S.A. 103: 5320–5325.

Dobin A, Davis CA, Schlesinger F, Drenkow J, Zaleski C, Jha S, Batut P, Chaisson M, Gingeras TR. 2013. STAR: ultrafast universal RNA-seq aligner. Bioinformatics 29: 15–21.

Drew HR, Travers AA. 1985. DNA bending and its relation to nucleosome positioning. J. Mol. Biol. 186: 773–790.

Duffy EE, Rutenberg-Schoenberg M, Stark CD, Kitchen RR, Gerstein MB, Simon MD. 2015. Tracking distinct RNA populations using efficient and reversible covalent chemistry. Mol. Cell 59: 858–866.

Fan X, Moqtaderi Z, Jin Y, Zhang Y, Liu XS, Struhl K. 2010. Nucleosome depletion in yeast terminator regions is not intrinsic and can occur by a transcriptional mechanism linked to 3’ end formation. Proc. Natl. Acad. Sci. U.S.A. 107: 17945–17950.

Fedor MJ, Lue NF, Kornberg RD. 1988. Statistical positioning of nucleosomes by specific protein binding to an upstream activating sequence in yeast. J. Mol. Bio. 204: 109–127.

Gibson DG, Benders GA, Axelrod KC, Zaveri J, Algire MA, Moodie M, Montague MG, Venter JC, Smith HO, Hutchison CA, 3rd. 2008. One-step assembly in yeast of 25 overlapping DNA fragments to form a complete synthetic Mycoplasma genitalium genome. Proc. Natl. Acad. Sci. U.S.A. 105: 20404–20409.

Gkikopoulos T, Schofield P, Singh V, Pinskaya M, Mellor J, Smolle M, Workman JL, Barton GJ, Owen-Hughes T. 2011. A role for Snf2-related nucleosome-spacing enzymes in genome-wide nucleosome organization. Science 333: 1758–1760.

Hahn S, Hoar ET, Guarente L. 1985. Each of three “TATA elements” specifies a subset of transcription initiation sites at the CYC1 promoter of Saccharomyces cerevisiae. Proc. Natl. Acad. Sci. U.S.A. 82: 8562–8566.

He F, Jacobson A. 2015. Nonsense-mediated mRNA decay: degradation of defective transcripts Is only part of the story. Annu. Rev. Genet. 49: 339–366.

Hughes A, Jin Y, Rando OJ, Struhl K. 2012. A functional evolutionary approach to identify determinants of nucleosome positioning: A unifying model for establishing the genome-wide pattern. Mol. Cell 48: 5–15.

Iyer V, Struhl K. 1995a. Mechanism of differential utilization of the his3 T_R_ and T_C_ TATA elements. Mol. Cell. Biol. 15: 7059–7066.

Iyer V, Struhl K. 1995b. Poly(dA:dT), a ubiquitous promoter element that stimulates transcription via its intrinsic structure. EMBO J. 14: 2570–2579.

Jin Y, Eser U, Struhl K, Churchman LS. 2017. The ground state and evolution of promoter regions directionality. Cell 170: 889–898.

Jin Y, Geisberg JV, Moqtaderi Z, Ji Z, Hoque M, Tian B, Struhl K. 2015. Mapping 3’ mRNA isoforms on a genomic scale. Curr. Protoc. Mol. Biol. 110: 4.23.21-24.23.17.

Joshi AA, Struhl K. 2005. Interaction of the Eaf3 chromodomain with methylated histone H3-K36 mediates preferential histone deacetylation at mRNA coding regions. Mol. Cell 20: 971–978.

Kaplan CD, Laprade L, Winston F. 2003. Transcription elongation factors repress transcription initiation from cryptic sites. Science 301: 1096–1099.

Keogh MC, Kurdistani SK, Morris SA, Ahn SH, Podolny V, Collins SR, Schuldiner M, Chin K, Punna T, Thompson NJ et al. 2005. Cotranscriptional Set2 methylation of histone h3 lysine 36 recruits a repressive rpd3 complex. Cell 123: 593–605.

Kim TK, Hemberg M, Gray JM, Costa AM, Bear DM, Wu J, Harmin DA, Laptewicz M, Barbara-Haley K, Kuersten S et al. 2010. Widespread transcription at neuronal activity-regulated enhancers. Nature 465: 182–187.

Kouprina N, Larionov V. 2008. Selective isolation of genomic loci from complex genomes by transformation-associated recombination cloning in the yeast Saccharomyces cerevisiae. Nat. Protoc. 3: 371–377.

Kourennaia OV, Tsujikawa L, deHaseth PL. 2005. Mutational analysis of Escherichia coli heat shock transcription factor Sigma 32 reveals similarities with Sigma 70 in recognition of the −35 promoter element and differences in promoter DNA melting and −10 recognition. J. Bact. 187: 6762–6769.

Krietenstein N, Wal M, Watanabe S, Park B, Peterson CL, Pugh BF, Korber P. 2016. Genomic nucleosome organization reconstituted with pure proteins. Cell 167: 709–721 e712.

Kristjuhan A, Svejstrup JQ. 2004. Evidence for distinct mechanisms facilitating transcript elongation through chromatin in vivo. EMBO J. 23: 4243–4252.

Kurosaki T, Popp MW, Maquat LE. 2019. Quality and quantity control of gene expression by nonsense-mediated mRNA decay. Nat. Rev. Mol. Cell Biol. 20: 406–420.

Langmead B, Salzberg SL. 2012. Fast gapped-read alignment with Bowtie 2. Nat. Methods 9: 357–359.

Lee W, Tillo D, Bray N, Morse RH, Davis RW, Hughes TR, Nislow C. 2007. A high-resolution atlas of nucleosome occupancy in yeast. Nat. Genet. 39: 1235–1244.

Lorch Y, Maier-Davis B, Kornberg RD. 2014. Role of DNA sequence in chromatin remodeling and the formation of nucleosome-free regions. Genes Dev. 28: 2492–2497.

Lui KH, Geisberg JV, Moqtaderi Z, Struhl K. 2022. 3’ untranslated regions are modular entities that determine polyadenylation profiles. Mol. Cell. Biol. 42: e0024422.

Machida RJ, Lin YY. 2014. Four methods of preparing mRNA 5’ end libraries using the Illumina sequencing platform. PLoS One 9: e101812.

Mason PB, Struhl K. 2003. The FACT complex travels with elongating RNA polymerase II and is important for the fidelity of transcriptional initiation in vivo. Mol. Cell. Biol. 23: 8323–8333.

Mavrich TN, Ioshikhes IP, Venters BJ, Jiang C, Tomsho LP, Qi J, Schuster SC, Albert I, Pugh BF. 2008. A barrier nucleosome model for statistical positioning of nucleosome throughout the yeast genome. Genome Res. 18: 1073–1083.

Miller C, Schwalb B, Maier K, Schulz D, Dumcke S, Zacher B, Mayer A, Sydow J, Marcinowski L, Dolken L et al. 2011. Dynamic transcriptome analysis measures rates of mRNA synthesis and decay in yeast. Mol. Syst. Biol. 7: 458.

Moqtaderi Z, Geisberg JV, Jin Y, Fan X, Struhl K. 2013. Species-specific factors mediate extensive heterogeneity of mRNA 3’ ends in yeasts. Proc. Natl. Acad. Sci. U.S.A. 110: 11073–11078.

Nagawa F, Fink GR. 1985. The relationship between the “TATA” sequence and transcription initiation sites at the HIS4 gene of Saccharomyces cerevisiae. Proc. Natl. Acad. Sci. U.S.A. 82: 8557–8561.

Neil H, Malabat C, d’Aubenton-Carafa Y, Xu Z, Steinmetz LM, Jacquier A. 2009. Widespread bidirectional promoters are the major source of cryptic transcripts in yeast. Nature 457: 1038–1052.

Pelechano V, Wei W, Steinmetz LM. 2013. Extensive transcriptional heterogeneity revealed by isoform profiling. Nature 497: 127–131.

Pelechano V, Wei W, Steinmetz LM. 2016. Genome-wide quantification of 5’-phosphorylated mRNA degradation intermediates for analysis of ribosome dynamics. Nat. Protoc. 11: 359–376.

Proudfoot NJ. 2011. Ending the message: poly(A) signals then and now. Genes Dev. 25: 1770–1782.

Proudfoot NJ, Brownlee GG. 1976. 3’ non-coding region sequences in eukaryotic messenger RNA. Nature 263: 211–214.

Quinlan AR, Hall IM. 2010. BEDTools: a flexible suite of utilities for comparing genomic features. Bioinformatics 26: 841–842.

Rougemaille M, Libri D. 2010. Control of cryptic transcription in eukaryotes. Adv. Exp. Med. Biol. 702: 122–131.

Schwabish MA, Struhl K. 2004. Evidence for eviction and rapid deposition of histones upon transcriptional elongation by RNA polymerase II. Mol. Cell. Biol. 24: 10111–10117.

Schwabish MA, Struhl K. 2006. Asf1 mediates histone eviction and deposition during elongation by RNA polymerase II. Mol. Cell 22: 415–422.

Schwabish MA, Struhl K. 2007. The Swi/Snf complex is important for histone eviction during transcriptional activation and RNA polymerase II elongation in vivo. Mol. Cell. Biol. 27: 6987–6995.

Schwanhausser B, Busse D, Li N, Dittmar G, Schuchhardt J, Wolf J, Chen W, Selbach M. 2011. Global quantification of mammalian gene expression control. Nature 473: 337–342.

Segal E, Widom J. 2009. Poly(dA:dT) tracts: major determinants of nucleosome organization. Curr. Opin. Struct. Biol. 19: 65–71.

Sekinger EA, Moqtaderi Z, Struhl K. 2005. Intrinsic histone-DNA interactions and low nucleosome density are important for preferential accessibility of promoter regions in yeast. Mol. Cell 18: 735–748.

Struhl K. 1985. Naturally occurring poly(dA-dT) sequences are upstream promoter elements for constitutive transcription in yeast. Proc. Natl. Acad. Sci. U.S.A. 82: 8419–8423.

Struhl K. 1986. Constitutive and inducible Saccharomyces cerevisiae promoters: evidence for two distinct molecular mechanisms. Mol. Cell. Biol. 6: 3847–3853.

Struhl K. 2007. Transcriptional noise and the fidelity of initiation by RNA polymerase II. Nat. Struct. Mol. Biol. 14: 103–105.

Struhl K, Segal E. 2013. Determinants of nucleosome positioning. Nat. Struct. Mol. Biol. 20: 267–273.

Sun M, Schwalb B, Pirkl N, Maier KC, Schenk A, Failmezger H, Tresch A, Cramer P. 2013. Global analysis of eukaryotic mRNA degradation reveals xrn1-dependent buffering of transcript levels. Mol. Cell 52: 52–62.

Thiebaut M, Kisseleva-Romanova E, Rougemaille M, Boulay J, Libri D. 2006. Transcription termination and nuclear degradation of cryptic unstable transcripts: a role for the nrd1-nab3 pathway in genome surveillance. Mol. Cell 23: 853–864.

Wickens M, Stephenson P. 1984. Role of the conserved AAUAAA sequence: four AAUAAA point mutants prevent messenger RNA 3’ end formation. Science 226: 1045–1051.

Winston F, Dollard C, Ricupero-Hovasse SL. 1995. Construction of a set of convenient Saccharomyces cerevisiae strains that are isogenic to S288C. Yeast 11: 53–55.

Wong K-H, Jin Y, Moqtaderi Z. 2013. Multiplex Illumina sequencing using DNA barcoding. Curr. Protoc. Mol. Biol. Chapter 7: Unit 7.11.

Wyers F, Rougemaille M, Badis G, Rousselle JC, Dufour ME, Boulay J, Regnault B, Devaux F, Namane A, Seraphin B et al. 2005. Cryptic Pol II transcripts are degraded by a nuclear quality control pathway involving a new poly(A) polymerase. Cell 121: 725–737.

Xu Z, Wei W, Gagneur J, Perocchi F, Clauder-Munster S, Camblong J, Guffanti E, Stutz F, Huber W, Steinmetz LM. 2009. Bidirectional promoters generate pervasive transcription in yeast. Nature 457: 1033–1037.

Yuan G-C, Liu Y-J, Dion MF, Slack MD, Wu LF, Altschuler SJ, Rando OJ. 2005. Genome-scale identification of nucleosome positions in S. cerevisiae. Science 309: 626–630.

Zhang Y, Moqtaderi Z, Rattner BP, Euskirchen G, Snyder M, Kadonaga JT, Liu XS, Struhl K. 2009. Intrinsic histone-DNA interactions are not the major determinant of nucleosome positions in vivo. Nat. Struct. Mol. Biol. 16: 847–852.

